# Urbanization drives parallel shifts in morphology and colouration of *Pieris rapae* across three cities

**DOI:** 10.64898/2026.02.11.705400

**Authors:** Catherine A. McManus, Matthew H. Koski, Rachel B. Spigler

**Affiliations:** Department of Biology, Temple University, Philadelphia, PA, USA; Department of Biological Sciences, Clemson University, SC, USA

**Keywords:** urban evolution, parallel evolution, phenotypic divergence, flight morphology, wing pigmentation, near-infrared reflectance, sexual dimorphism, habitat fragmentation, urban heat island, *Pieris rapae*

## Abstract

Urbanization is a major driver of global environmental change, yet the evolutionary consequences for natural populations remain poorly understood. The cabbage white butterfly, *Pieris rapae*, is one of few butterfly species that thrives in cities, offering a unique opportunity to investigate whether urbanization drives parallel phenotypic evolution. In this study, we quantified phenotypic divergence between urban and non-urban populations across three independent metropolitan regions (Cleveland, Philadelphia, and Pittsburgh, U.S.), measuring body size, flight morphology, and wing pigmentation and reflectance in >500 field-caught individuals from 19 sites. Across regions, we found repeated reductions in overall body size and relative thorax mass in urban butterflies of both sexes, along with reduced relative forewing area in urban males, suggesting sex-specific impacts on mobility. Wing pigmentation showed limited urban-associated variation, but we detected novel sexual dimorphism in near-infrared reflectance within urban populations. Together, these results demonstrate repeatable divergence between urban and non-urban butterfly populations across multiple trait axes and suggest that plastic responses to urban stressors previously documented in experimental work manifest as consistent phenotypic shifts in natural populations.

## Background

Urban land cover has expanded dramatically over recent centuries and continues to grow at unprecedented rates and scales [1]. The resulting environmental challenges are severe: human infrastructure and habitat loss subdivide populations while imposing intense selection pressures associated with greater impervious surface cover, elevated temperatures, increased pollution, and altered species interactions [2,3]. For relatively few species, preadaption to the strong environmental filters imposed by urban environments may secure their persistence [4]. For the remaining species, however, populations can only self-sustain via adaptive plasticity or rapid evolution. In these cases, we expect to see phenotypic differences between urban and non-urban conspecifics in traits relevant to urban environmental pressures. Indeed, studies comparing urban and non-urban populations of plants [5,6], invertebrates [7,8], reptiles [9], and mammals [10] have revealed stark phenotypic shifts. Most animal studies, however, focus on birds and mammals, with comparatively little work on insects [11]. Given the key roles insects play in ecosystem function and predictions that global urban land cover will increase to over 1.1 million km^2^ by 2100 [12], there is an urgency to understand ecological and evolutionary responses of insect populations to urbanization.

One key trait expected to differ between urban and non-urban populations is body size. Urban areas are significantly warmer than surrounding non-urban areas, forming urban heat islands (UHIs), with cities on average 2.9°C warmer and up to 9°C warmer than nearby non-urban areas [13]. Warmer temperatures can directly impact body size in insects by altering metabolic processes and development. Individuals developing in warmer temperatures tend to mature at smaller sizes, a pattern known as the temperature-size rule [14–16]. Beyond these plastic responses to temperature, urban environments may also select for smaller body size if reduced size confers energetic advantages. Because metabolic costs increase at higher temperatures in ectotherms [17] and smaller individuals require less absolute energy [18], smaller body size may be favored in resource-limited urban environments.

Habitat heterogeneity may also drive divergence in body size, as well as other aspects of flight morphology [19]. The need to locate patchily distributed floral resources and mates, as well as navigate complex urban landscapes, may favor phenotypes balancing speed, endurance, and manoeuvrability. For many flying insects, larger body size enables faster flight [20]. Additionally, larger thoraxes (which contain greater flight muscle mass) and larger wings enhance acceleration [21] and the ability to traverse longer distances [22]. Wing loading (body mass relative to wing area) and aspect ratio (wing length relative to width) both influence manoeuvrability, with lower values conferring improved turning ability [23]. Thus, urban populations may increase in body, thorax, and wing size, countering expected reductions from temperature [24], while exhibiting lower wing loading and aspect ratios to enhance manoeuvrability.

Urban and non-urban insect populations may also differ in colouration because of its impact on thermoregulation. The thermal melanism hypothesis [25] predicts that warmer climates select against highly pigmented, less reflective individuals because they absorb more solar radiation and are at higher risk of overheating than lighter individuals. This hypothesis should also extend to the UHI effect, such that urban populations show overall reduced pigmentation or increased reflectance arising from evolutionary change [26], phenotypic plasticity (e.g., in response to temperature: [27]), or a combination of both [28]. However, spatial patterns of pigmentation and reflectance changes depend on how colouration interacts with thermoregulatory behaviour, such as which body surfaces are exposed to solar radiation during basking [29]. Studies on urban-non-urban divergence have not considered reflectance in the near-infrared (NIR) region, yet NIR is responsible for approximately 55% of solar energy reaching Earth [30]. Moreover, NIR reflectance responds more readily to changes in the thermal environment compared to visible colouration, as the latter is constrained by roles in camouflage and communication [30,31]. Decreasing pigmentation with increasing temperature has been demonstrated in insect studies conducted across large geographic scales, including elevational [32] or latitudinal clines [33], and over temporal scales related to climate change [34]. However, the thermal melanism hypothesis has only recently been applied to UHIs [35–38], and no urban studies have yet examined NIR reflectance.

Stressors are shared among cities, highlighting how cities have become ecologically similar [39–41]. Despite this, there is still notable environmental heterogeneity among cities in age, size, and climate [42] that may interact with the effects of urbanization to shape phenotypic variation. While most studies focus on a single urban center, those spanning multiple cities find variation in responses across cities [43,44], reinforcing the importance of replicating urban studies to assess the generality of responses. The absence of consistent parallelism is also a reminder that phenotypic differences among populations can arise from neutral processes. Cities can present sizeable barriers to gene flow [45], creating small, isolated populations where genetic drift and inbreeding can overwhelm natural selection.

Here, we explore whether differences in morphological traits related to body size and colouration are associated with urbanization and key environmental drivers in *Pieris rapae* (Lepidoptera: Pieridae), the cabbage white butterfly, across three metropolitan regions. Unlike many butterfly species, which tend to avoid urban areas, *P. rapae* exhibits a strong affinity for urban habitats and reaches high abundances in cities [46,47]. This urban success suggests that *P. rapae* may exhibit phenotypic plasticity or adaptive evolutionary responses that facilitate its survival and reproduction within anthropogenic landscapes.

We ask the following in *P. rapae*: 1) Is urbanization associated with shifts in body size traits, consistent with thermal or mobility pressures? 2) Is urbanization associated with changes in wing colouration (i.e. pigmentation and reflectance) predicted to mitigate thermal stress? 3) Are patterns consistent across geographic regions?

## Methods

### Study sites and sampling

We collected adult *P. rapae* butterflies (species details in electronic supplementary material, S1) in three metropolitan regions in the U.S. that vary in population size and density, climate, and intensity of urbanization: Philadelphia, PA, Pittsburgh, PA, and Cleveland, OH (figure 1, city descriptions in electronic supplementary material, S2.1). In each region, we identified three urban sites (with one additional urban site in Philadelphia) and three non-urban sites (additional site selection information in electronic supplementary material, S2.2 and table S1).

**Figure 1.**
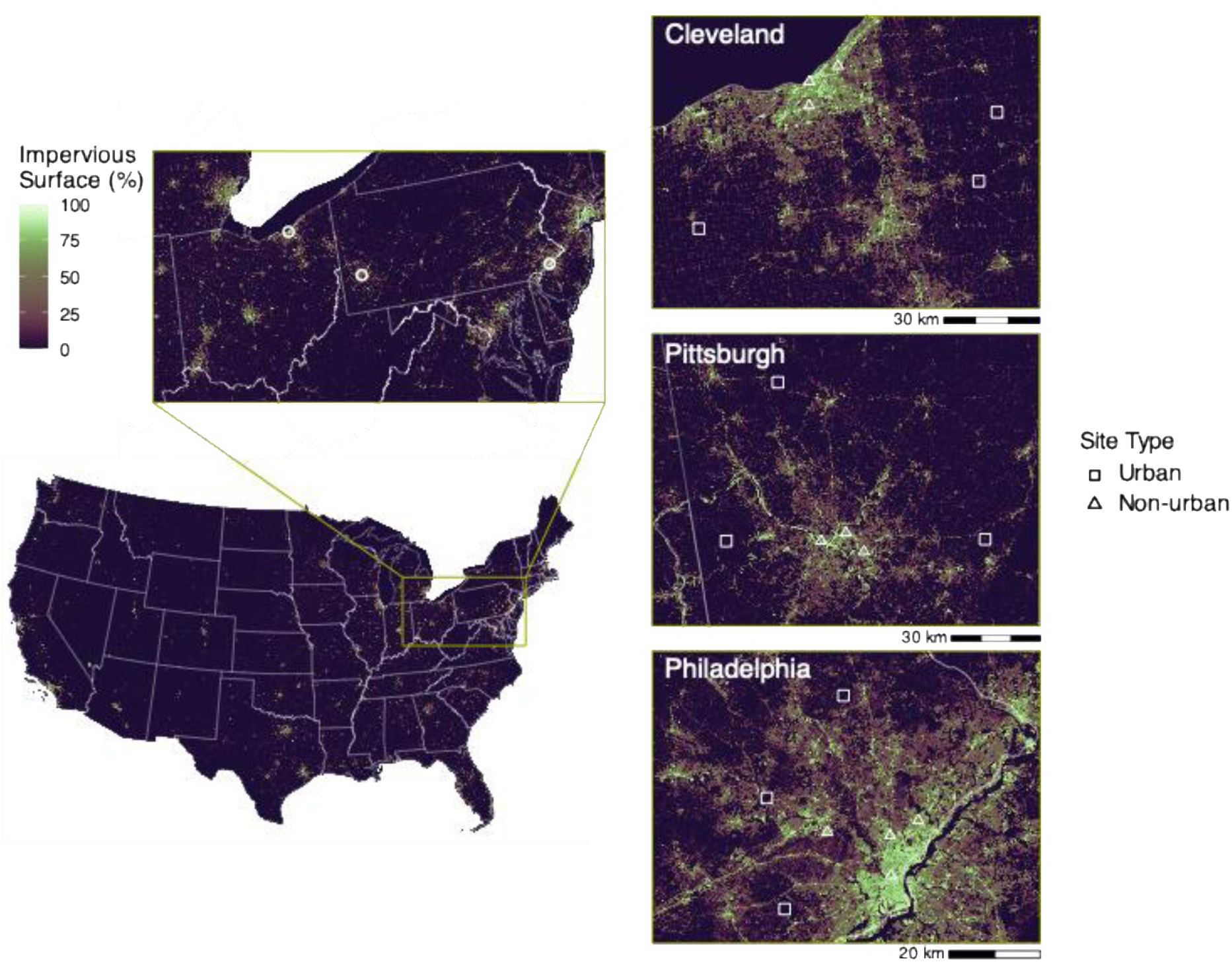
Locations of *Pieris rapae* urban (▪) and non-urban (▲) sampling sites across Cleveland, Pittsburgh, and Philadelphia. Background shows 2000–2022 mean impervious surface cover from the National Land Cover Database.

Collections took place from 16 June to 18 August 2022. Because *P. rapae* exhibits distinct spring/fall and summer forms that differ in pigmentation [48] and flight morphology [49,50], collections targeted the summer form to isolate the effects of urbanization from seasonal variation. We captured butterflies with hand nets and stored euthanized butterflies at −20°C. We collected 571 butterflies (409 males, 162 females) across the three regions. This male sampling bias is typical of *P. rapae* studies [51].

### Site Data

In addition to categorizing sites as urban or non-urban, we considered that stressors associated with urbanization can vary along a gradient. We quantified impervious surface cover, mean and maximum summer temperature, and patch complexity (perimeter area ratio, PARA) for each site using long term (2000-2022) averages (description of datasets and methods in electronic supplementary material, S2.3). Site-level data were calculated from 1 km circular buffers (for buffer size rationale, see electronic supplementary material, S2.4 and table S2). See electronic supplementary material, table S4 for site data.

After environmental variable reduction (electronic supplementary material, S2.5), we used maximum air temperature (hereafter, “temperature”) as our representative variable of the urban gradient. Because of the high collinearity among environmental variables in our study (electronic supplementary material, table S3), we cannot estimate the unique effects of temperature alone and recognize that observed patterns may reflect temperature, habitat heterogeneity, unmeasured covariates, or some combination of these factors.

### Morphological Data

All wing measurements were taken on the right side. The left wing was substituted only when the right wing was damaged or showed scale loss. We excluded butterflies with damage or scale loss on both wings.

#### Size morphometrics

Butterflies were thawed, placed in a humid container for >24 hours to relax, then pinned and spread. Pinned butterflies were stored at room temperature with desiccant and dried to constant mass. We used the final, constant mass (total mass) as our measure of overall body size. We subsequently dissected butterflies and measured thorax mass. Wings were photographed on dorsal and ventral sides following a standardized protocol (electronic supplementary material, S3.1).

We collected data on wing size from the digital images using ImageJ [52]. For each butterfly, the length of a single forewing was measured from the basal wing area to the wing tip. We measured the forewing area as the outline of the entire wing. We also calculated wing loading and aspect ratio, metrics associated with manoeuvrability. We calculated wing loading as total mass divided by total forewing area, following [21], with total forewing area estimated by doubling the single wing measurement. We calculated aspect ratio as 4 x (forewing length)^2^/ forewing area, following [21].

#### Pigmentation

The dorsal forewing of *P. rapae* is pigmented at the forewing tip and eyespots (one in males, two in females) (electronic supplementary material, figure S1a). We quantified the size of these areas following [48,53]. In ImageJ, we removed the image background, converted to grayscale (8-bit), and thresholded at 0 and 135 (where 0 = black, 255 = white). Pixels within this threshold were considered pigmented, and the area of these pixels was measured.

#### Reflectance

We used a reflectance spectrometer to record wing reflectance for 10 males and 10 females per site (or all available for sites with fewer than 10 per sex). We captured reflectance spectra at select wing spots relevant for thermoregulation in this species (electronic supplementary material, figure S1b; [54–56]) from 250-950 nm using a UV-NIR spectrometer (protocol in electronic supplementary material, S3.2). We calculated the average percent reflectance over the visible (400-700 nm), and near infrared (NIR, 700-950 nm) portions of the spectrum for each wing spot (hereafter, NIR-r and Vis-r).

### Statistical analyses

#### Correlations and variable reduction

We calculated pairwise Pearson correlation coefficients (*r*) to identify highly correlated traits (electronic supplementary material, figures S2 and S3). After variable reduction (electronic supplementary material, S4.1), we retained thirteen traits for analysis: four morphometric traits (total mass, thorax mass, forewing area, and aspect ratio), two pigmentation traits (wing tip area and anterior wing spot area), and seven reflectance traits (postdiscal NIR-r and Vis-r, wing tip NIR-r, basal forewing NIR-r, basal hindwing dorsal NIR-r, basal hindwing ventral NIR-r, and marginal hindwing NIR-r). Sample sizes for each trait varied (electronic supplementary material, table S5).

#### Testing impacts of urbanization

We first tested whether urban and non-urban sites differed in impervious surface cover, temperature, and habitat complexity (PARA). These tests assessed how well the categorical designations captured the underlying environmental gradients and identified regional differences. We tested for main effects of urbanization classification (urban vs non-urban), region, and their interaction.

We used linear mixed-effects models (Gaussian distribution) to test the effects of urbanization on morphological traits. For each trait, we fit two models: one treating urbanization as categorical (urban vs non-urban; hereafter U-NU) and another using mean-centered temperature as a continuous abiotic gradient associated with urbanization. All models included fixed effects of metropolitan region (hereafter, “region”) and sex to account for overall regional differences and sexual dimorphism, respectively. We included an interaction between our main predictor variable (U-NU or temperature) and region. A significant interaction would indicate that responses to urbanization differed among regions. We also included an interaction between the main predictor and sex to account for potential sex-specific responses. Site was treated as a random effect to account for non-independence among samples from the same site. For thorax mass and forewing area, we included body mass as a covariate to account for allometric scaling and evaluate changes beyond overall body size differences. We similarly included total wing area as a covariate in our models of wing pigmentation area.

Because wing spot pigmentation and reflectance influence body temperature differently across wing areas due to interactions with basking postures in *P. rapae* [54–56], we analyzed all traits separately. During reflectance basking, distal wing regions direct more solar energy toward the body when reflectance is high [54–56]. Therefore, contrary to the typical expectation that warmer temperatures favor reduced pigmentation and increased reflectance, predictions for distal wing areas are reversed. We predicted that urban butterflies would show increased pigmentation (larger wing tip and wing spot) and reduced reflectance (lower postdiscal NIR-r and Vis-r and wing tip NIR-r) in these areas to limit heat gain during reflectance basking (electronic supplementary material, figure S1b). In contrast, the basal forewing, basal hindwing (dorsal and ventral), and marginal hindwing absorb radiation during dorsal and ventral basking [54–56]. We therefore predicted increased reflectance (higher NIR-r) in these areas to reduce heat absorption (electronic supplementary material, figure S1b), aligning with standard thermal melanism expectations.

Models were fit with nlme [57,58] (full description of model testing procedure in electronic supplementary material, S4.2). To address the inflated risk of type I errors from testing multiple traits [59], we controlled the false discovery rate (FDR) by adjusting *p* values for each of the three families of hypotheses (morphometrics, pigmentation, and reflectance) using the Benjamini-Hochberg procedure [60]. We used emmeans [61] to estimate marginal means, predict slopes for the effects of temperature, and perform comparisons between urban and non-urban butterflies, between sexes, and among regions (with Tukey-adjusted *p* values for the three pairwise region comparisons). We report and plot marginal means ± standard error and estimated slopes with 95% confidence intervals. For each trait, we compared the fit of the U-NU model against the fit of the temperature model using AIC. We considered a reduction in AIC >2 as evidence of a better model fit. We plotted model estimates and raw data using ggplot2 (v. 3.5.1) [62]. All data analyses and visualizations were performed in R (v. 4.4.2) [63].

## Results

### Site data

Urban and non-urban sites showed pronounced differences across environmental metrics of urbanization. Because we classified sites using a 50% impervious surface threshold, differences in impervious surface were expected. Nonetheless, the realized difference was striking, with over 20 times greater impervious surface cover in urban sites compared to non-urban sites (*F*_1_ = 675.8, *p* < 0.001, figure 2a). Urban sites also displayed over 16 times more habitat complexity than non-urban sites, based on PARA (*F*_1_ = 27.2, *p* < 0.001, figure 2b). Mean impervious surface cover and PARA were similar across regions (cover: *F*_2_ = 3.3, *p* = 0.07; PARA: *F*_2_ = 0.32, *p* = 0.73), and there was no interaction between site type and region (cover: *F*_2_ = 0.21, *p* = 0.82; PARA: *F*_2_ = 0.17, *p* = 0.84).

**Figure 2.**
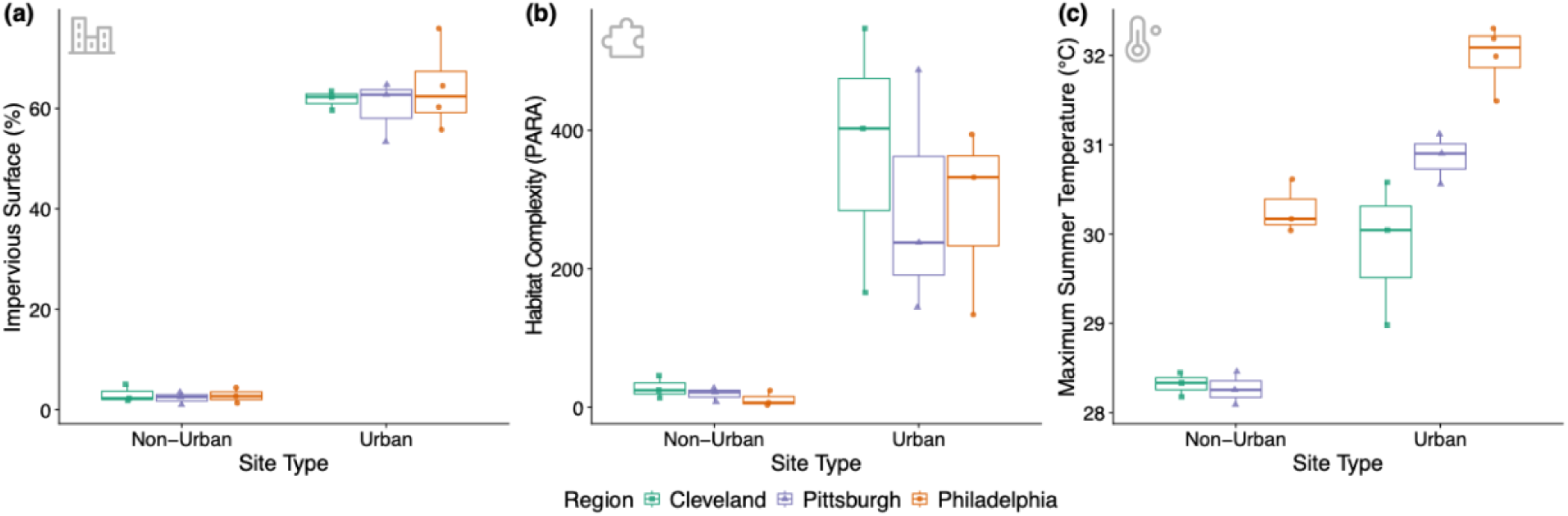
Environmental characteristics of non-urban and urban sampling sites across three metropolitan regions. (a) Impervious surface cover, (b) habitat complexity (perimeter to area ratio of undeveloped patches), and (c) maximum summer air temperature. Points represent sampling sites (Cleveland *n* = 6, Pittsburgh *n* = 6, Philadelphia *n* = 7).

In contrast, our sites spanned a gradient of temperatures (figure 2c). Still, urban sites were on average 2.0 ± 0.19°C warmer than non-urban sites based on average maximum summer temperature (*F*_1_ = 106.9, *p* < 0.001). There were also differences in temperature across regions. Philadelphia was 2.0 ± 0.23°C warmer than Cleveland (*t*_13_ = 8.9, *p* < 0.001) and 1.6 ± 0.23°C warmer than Pittsburgh (*t*_13_ = 6.9, *p* < 0.001), which was similar in magnitude to the difference between urban and non-urban sites. Mean temperatures of Cleveland and Pittsburgh did not differ (*t*_13_ = 2.0, *p* = 0.15), and there was no interaction between site type and region (*F*_2_ = 2.9, *p* = 0.09).

### Size morphometrics

*Pieris rapae* from urban sites were significantly lighter in total body mass than those from non-urban sites (figure 3a, electronic supplementary material, table S6). On average, urban butterflies were 1.9 ± 0.47 mg smaller, a 10.2% reduction. Prior to controlling for FDR, model results showed an interaction between U-NU and sex, driven by significant mass reductions in males (*t*_13_ = 5.2, *p* < 0.001) but not females (*t*_13_ = 1.7, *p* = 0.11). The impact of urbanization was roughly equivalent to regional differences; compared to Philadelphia, butterflies from Cleveland were 2.0 ± 0.55 mg lighter (*t*_13_ = 3.7, *p* = 0.007), representing a 10.4% decrease, and those from Pittsburgh were 2.3 ± 0.52 mg lighter (*t*_13_ = 4.5, *p* = 0.002), corresponding to a 12% decrease. There was no difference in overall size between butterflies from Cleveland and Pittsburgh (*t*_13_ = 0.56, *p* = 0.84). The interaction between U-NU and region was not significant, and mean total mass was similar between sexes. Using temperature as a continuous predictor variable yielded the same qualitative results: total mass decreased with increasing temperature (*β* = −0.97 mg/°C, CI: −1.42 to - 0.52; figure 3b, electronic supplementary material, table S6).

**Figure 3.**
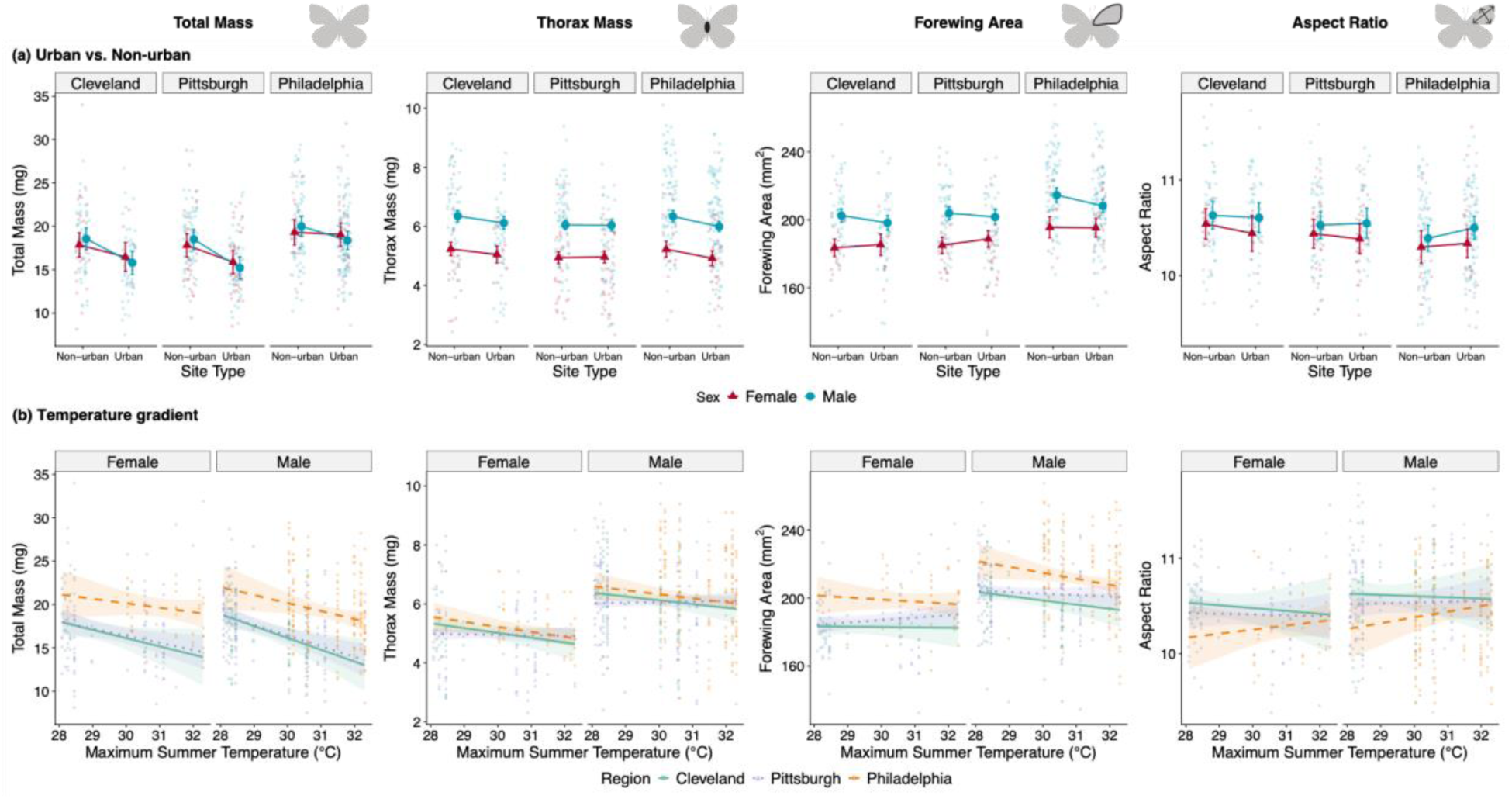
Effects of urbanization and temperature on size and mobility traits of *Pieris rapae*. (a) Urban vs non-urban categorical model, shown separately for each metropolitan region. (b) Temperature gradient (pooled across regions), shown separately for each sex.

Urban butterflies also had lower thorax mass than non-urban butterflies, even after adjusting for total mass (figure 3a, electronic supplementary material, table S6). Thorax mass of urban butterflies was 0.18 ± 0.08 mg, or 3%, less compared to non-urban butterflies for a given total mass, though evidence for this effect weakened after FDR correction. The effect of temperature was robust to correction for multiple tests: thorax mass declined 0.10 mg per °C (CI: −0.18 to −0.02; figure 3b, electronic supplementary material, table S6). There was no significant interaction between U-NU and region; there was, however, suggestive evidence of an interaction between temperature and region (*p* < 0.10 prior to FDR correction). Region-specific slopes revealed declines in Philadelphia (*β* = −0.15 mg/°C, CI: −0.30 to −0.01) and Cleveland (*β* = −0.15 mg/°C, CI: −0.29 to 0), but not Pittsburgh (*β* = 0 mg/°C, CI: −0.08 to 0.09). We also detected sexual dimorphisms: males had 1.1 ± 0.07 mg, or 21.5%, heavier thoraxes than females for a given total mass.

Neither urbanization category nor temperature had overall effects on forewing area, but there was suggestive evidence of sex-specific effects for each predictor (*p* < 0.10 prior to FDR correction; figure 3; electronic supplementary material, table S6). Although the interactions did not reach statistical significance, contrasts supported divergent responses between males and females. In the U-NU model, urban males had a small but significant reduction in forewing area compared to non-urban males of 4.3 ± 1.8 mm^2^, equivalent to ∼2% (*t*_13_ = 2.4, *p* = 0.03), but there was no difference between urban and non-urban females (*t*_13_ = 0.62, *p* = 0.55). Similarly, male forewing area declined with temperature (*β* = −2.3 mm^2^/°C, CI: −4.1 to −0.53), but female forewing size did not (*β* = 0.001 mm^2^/°C, CI: −2.4 to 2.4). On average, male wings were larger than female wings, with sex differences larger than U-NU differences. Male forewings were 15.9 mm^2^ (8.5%) larger than female forewings. There were regional differences with Philadelphia butterflies having larger wings (203 ± 1.43 mm^2^) than those from Cleveland (192 ± 1.40 mm^2^; *t*_13_ = 5.8, *p* < 0.001) and Pittsburgh (195 ± 1.22 mm^2^; *t*_13_ = 4.6, *p* = 0.001). There was no difference in wing size between Cleveland and Pittsburgh butterflies (*t*_13_ = 1.3, *p* = 0.40), and there were no interactions between region and our metrics of urbanization.

Aspect ratio did not vary with urbanization or temperature (figure 3, electronic supplementary material, table S6). There was, however, an effect of sex: males had ∼1% higher aspect ratios than females. We did not detect regional differences nor any significant interaction terms.

### Pigmentation

Wing tip area tended to increase in urban butterflies by 1.0 ± 0.50 mm^2^, or ∼8%, compared to non-urban butterflies and with temperature (*β* = 0.43 mm^2^/°C, CI: −0.10 to 0.95; figure 4; electronic supplementary material, table S7). These effects approached significance, but only prior to FDR correction. There were stronger regional differences: Philadelphia butterflies had smaller wing tips than those from Cleveland (2.8 ± 0.62 mm^2^ difference, *t*_13_ = 4.5, *p* = 0.002) and Pittsburgh (2.7 ± 0.56 mm² difference, *t*_13_ = 4.8, *p* = 0.001). Both differences represent roughly 20% reductions for butterflies from Philadelphia. There were no differences in wing tip size between Cleveland and Pittsburgh butterflies (*t*_13_ = 0.14, *p* = 0.99). Female wing tips were 3.8 ± 0.44 mm^2^ larger on average than those of males, a 34% increase. No interaction terms were significant.

**Figure 4.**
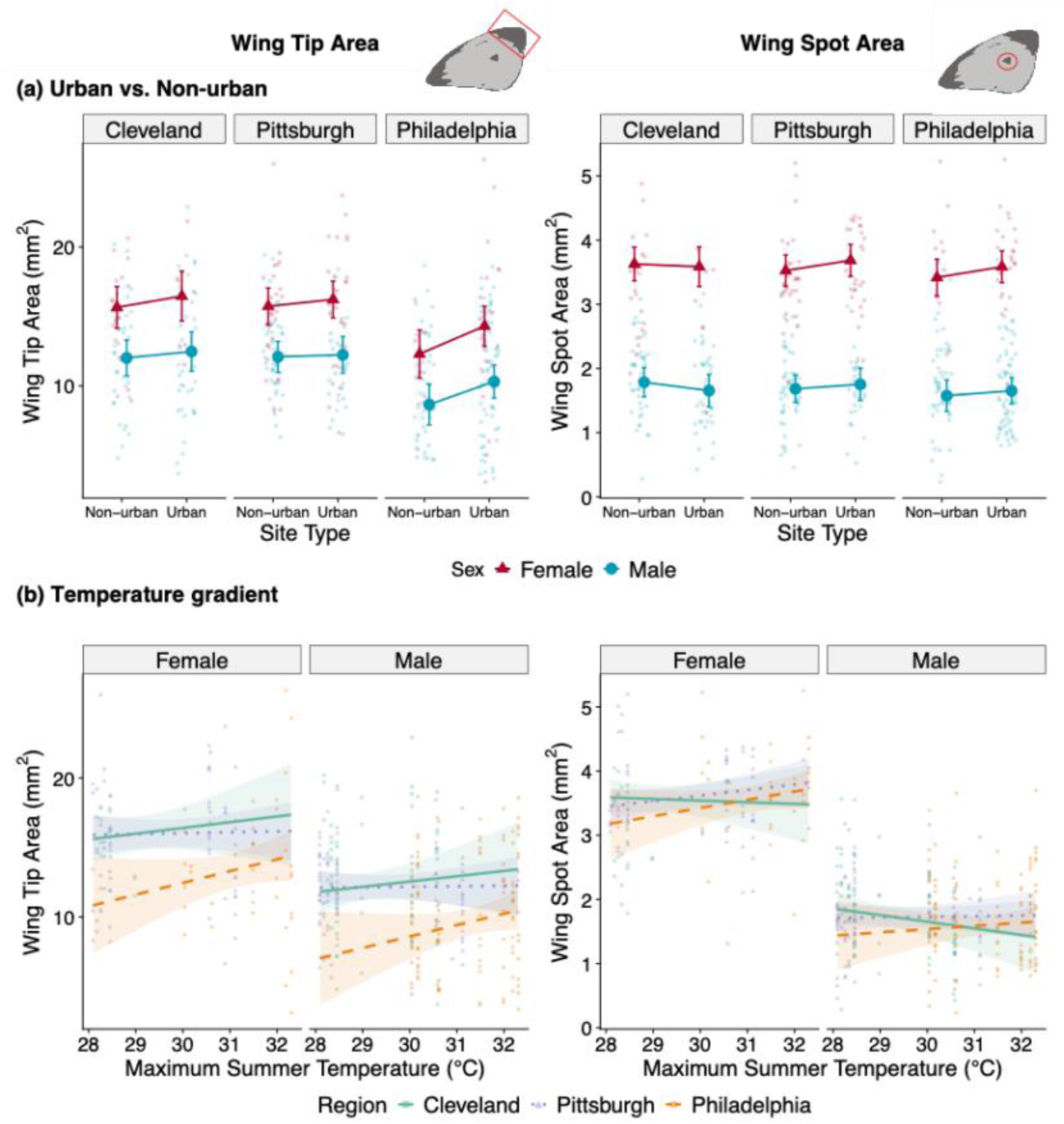
Effects of urbanization and temperature on wing pigmentation of *Pieris rapae*. (a) Urban vs nonurban effects, shown separately for each region. (b) Continuous temperature gradient, shown separately for each sex.

Posterior wing spot area did not vary with urbanization, temperature, or region (figure 4, electronic supplementary material, table S7). Although the interaction between temperature and sex approached significance before FDR correction, neither sex-specific slope significantly differed from zero (males: *β* = −0.01 mm^2^/°C, CI: −0.11 to 0.08; females: *β* = 0.06 mm^2^/°C, CI: −0.05 to 0.17). Females had wing spots twice as large as males (3.6 ± 0.07 mm^2^ vs 1.7 ± 0.05 mm^2^). No other model effects were significant.

### Reflectance

We detected a significant interaction between U-NU categorization and sex for postdiscal NIR reflectance prior to controlling for FDR (figure 5a; electronic supplementary material, table S8). Contrasts of sex-specific responses indicate that females showed a slight increase in percent reflectance in urban compared to non-urban butterflies (63.1 ± 1.2% vs 60.3 ± 1.2%), while reflectance slightly decreased in urban compared to non-urban males (56.4 ± 0.90% vs 58.6 ± 0.87%). While neither shift was significant (males: *t*_13_ = 1.8, *p* = 0.10; females: *t*_13_ = 1.7, *p* = 0.12), together the opposing weak patterns were sufficient to result in sexual dimorphism in urban sites. Using post hoc comparisons, we found that postdiscal NIR-r was similar between males and females within non-urban sites (*t*_284_ = 1.2, *p* = 0.24) but differed between males and females within urban sites (*t*_284_ = 4.6, *p* < 0.001) where females reflected 6.6 ± 1.5% more NIR.

**Figure 5.**
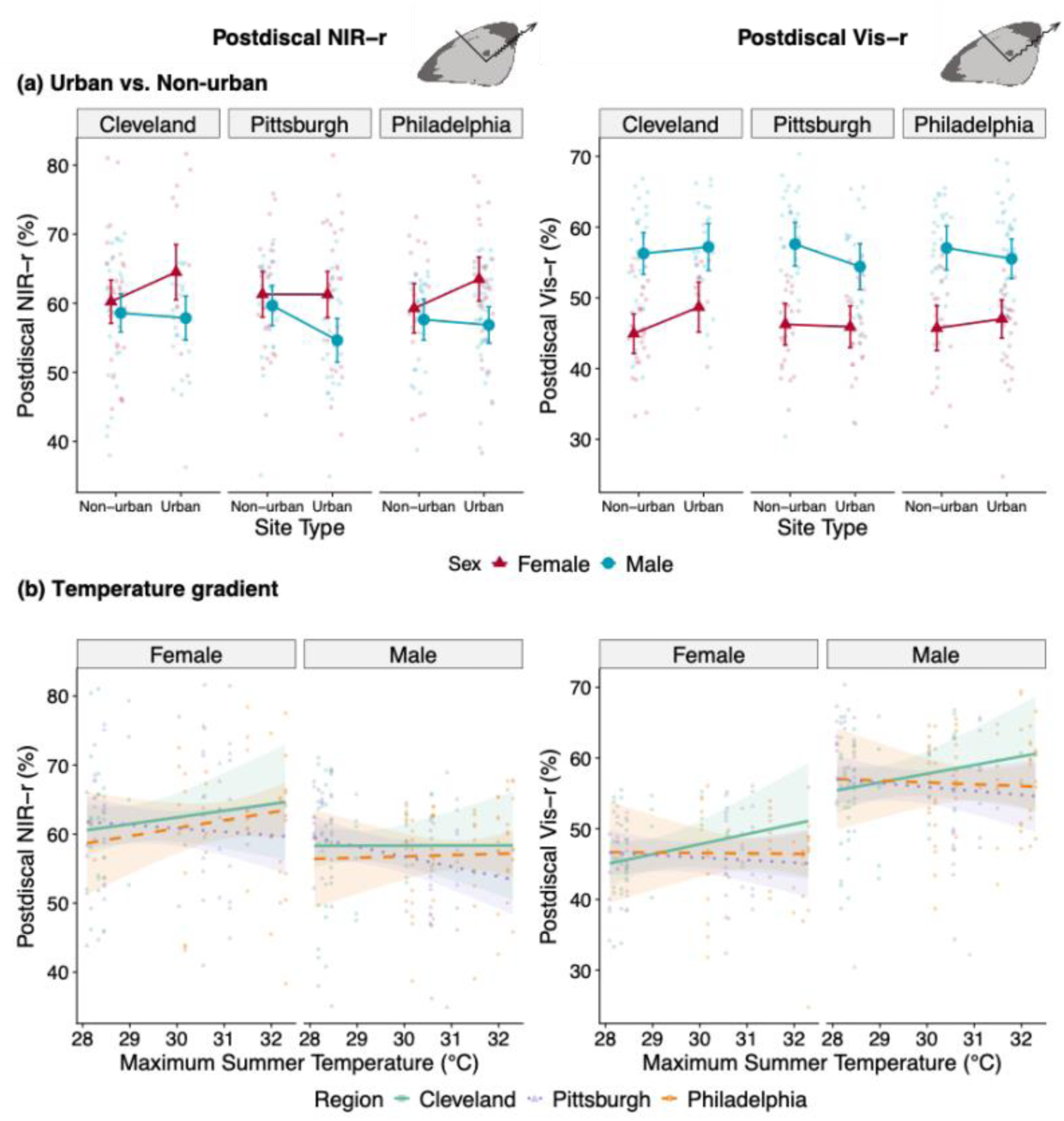
Effects of urbanization and temperature on visible reflectance (Vis-r) and near-infrared reflectance (NIR-r) at the postdiscal wing region of *Pieris rapae*. (a) Urban vs non-urban effects, shown separately for each region. (b) Continuous temperature gradient, shown separately for each sex.

We found a similar pattern for postdiscal visible reflectance: a U-NU by sex interaction that was marginally significant before FDR correction (figure 5a, electronic supplementary material, table S8). Again, this pattern was due to weak trends of higher Vis-r in urban females compared to non-urban females (47.2 ± 0.92% vs 45.6 ± 0.90%) and lower Vis-r in urban males compared to non-urban males (55.7 ± 0.97% vs 57.0 ± 0.95%). In contrast to NIR-r, however, sexual dimorphism was evident across site types and in the opposite direction. On average across both site types, males had ∼10% higher postdiscal Vis-r than females. Temperature did not explain variation in postdiscal NIR-r or VIS-r (figure 5b, electronic supplementary material, table S8).

Neither U-NU nor temperature significantly affected reflectance across the remaining wing areas examined (electronic supplementary material, table S8 and figure S4). We detected sexual dimorphism at the forewing tip with females showing 4.4 ± 0.85% greater NIR-r than males. We did not detect regional differences in reflectance in any wing area.

### Model selection

The U-NU model provided a better fit than the temperature model (ΔAIC > 2) for all traits (electronic supplementary material, tables S6-S8), indicating that urbanization effects were better captured by discrete urban vs non-urban classifications rather than continuous temperature gradients.

## Discussion

We present strong evidence of repeated morphometric divergence between urban and non-urban *P. rapae* populations across three metropolitan regions, demonstrating that urbanization drives parallel phenotypic shifts. These consistent patterns across independent cities suggest that urban environments impose predictable selection pressures or elicit repeated plastic responses. We further demonstrated associations between these traits and temperature, consistent with UHI effects. For all traits, however, a categorical urban-non-urban classification outperformed temperature alone, suggesting that multiple aspects of urbanization (e.g., impervious surfaces, pollution, and UHI) jointly influence these traits. In contrast, colouration was largely insensitive to urbanization, although we observed novel sexual dimorphism in NIR reflectance in urban environments, pointing to potential avenues for further study.

### Reduced body size in urban *P. rapae*

Urban *P. rapae* were consistently smaller than non-urban conspecifics across the three metropolitan regions in our study. Specifically, we observed decreases in both total mass and relative thorax mass in urban populations. Variation in these traits was negatively associated with temperature, consistent with expectations from the temperature-size rule [14] applied to UHI effects. Post hoc comparisons indicated that these patterns further extend to forewing size in males but not females. Although our results do not allow direct inference about the mechanism underlying this divergence, evidence from previous studies suggests phenotypic plasticity plays a primary role. Multiple experimental studies have documented that *P. rapae* reared under warmer temperatures mature at smaller sizes [64–69], and recent common garden experiments using Cleveland populations found no genetically based size differences between urban and non-urban butterflies [69]. These results suggest that the body size differences in our field study reflect plastic responses to developmental temperature. Our study complements experimental work by demonstrating that laboratory-documented plasticity manifests as consistent size variation across urban and non-urban populations.

The repeated urban-non-urban size differences across all three metropolitan regions argue against drift as an explanation, contrasting with size increases documented in urban *P. rapae* from Marseille, France that were attributed to drift [51]. While alternative processes such as inbreeding depression could also contribute to smaller body sizes [70–72], preliminary genomic analyses do not indicate elevated inbreeding in the urban populations in this study (Authors, unpublished data). Overall, the consistency of size reductions with urbanization and their association with temperature suggest an important role of environmental effects, though disentangling plastic and evolutionary responses will require additional common garden experiments across all three regions.

Interestingly, regional differences in body and wing size were comparable in magnitude to differences generated by urbanization, yet they did not align with expectations based on regional temperature differences. Butterflies from Philadelphia, the warmest region in our study, were the largest, contradicting the temperature-size rule and the within-region urban reductions we observed. This pattern mirrors results from a broader geographical analysis of *P. rapae.* Kingsolver et al. [67] found that populations from warmer southern U.S. regions were larger than those from cooler northern U.S. regions.

Together, these patterns suggest that fine-scale urban thermal environments are not equivalent to regional climate variation, potentially due to differences in spatial scale and temporal variability [73]. Therefore, the consequences of reduced body size in urban populations may not parallel those associated with large-scale climatic gradients.

Changes in body size have important fitness implications in insects [74,75]. Female size is positively correlated with fecundity in *P. rapae* [66,76], raising the possibility that smaller urban females exhibit decreased reproductive output. At the same time, smaller body size is associated with faster development and earlier reproduction in this species [77], highlighting life history trade-offs that may shape reproductive success in urban environments. For multivoltine species in particular, reduced generation time associated with smaller body size could allow additional generations per season, increasing population growth, as documented in urban populations of water fleas [78]. Such a mechanism is plausible in *P. rapae*, given its genetic variation in pupal size, developmental rate, and relative growth [77] and latitudinal and climatic gradients of development time and generation number [68,79]. Understanding whether urbanization-driven size reduction enables faster generation times could explain *P. rapae*’s remarkable persistence across urban landscapes.

### Sex-specific shifts in flight morphology in urban *P. rapae*

While body size is widely studied in insects due to its central role in fitness, few studies have evaluated urban-non-urban divergence in flight-related traits alongside overall body size (but see: [80,81]). Thorax mass directly reflects investment in flight muscles, and both thorax mass and wing size positively predict flight speed in *P. rapae* and other butterflies [21,50,65]. In our study, we found sex-specific responses to urbanization. Urban butterflies of both sexes showed reduced relative thorax mass and only males showed reduced wing area, morphological changes that together suggest reduced flight capacity in urban populations, with potentially stronger effects in males. Although our smaller female sample sizes could affect this pattern, experimental evidence supports sex-specific plasticity in wing size of *P. rapae*, with males showing steeper reaction norms than females [69]. Given that female natal dispersal is critical for genetic connectivity in fragmented landscapes [82], future work should investigate whether sex-specific selection pressures maintain female wing size relative to males despite urbanization effects.

Wing loading (body mass/wing area) is typically used as an estimate of flight performance and manoeuvrability in butterflies [23]. However, because it was nearly perfectly correlated with body mass in our dataset (*r* = ∼0.9), we did not present the decreased wing loading observed in urban populations. One interpretation of this result is that body mass declined disproportionately relative to wing area in urban butterflies, as documented in urban bees [81], which could lower wing loading and enhance flight [65,83]. However, because of the strong collinearity with body size, we cannot draw strong conclusions on wing loading alone. In addition, we did not find changes in wing aspect ratio (shape) associated with urbanization, though aspect ratio is a simplified metric of wing shape [23], and more detailed geometric morphometric analyses may reveal subtle shape changes not captured by our measurements. Instead, our data on thorax mass and wing size, which showed significant effects even after adjusting for body size, provide stronger evidence for urban-associated shifts in flight morphology.

### Limited changes in pigmentation and reflectance associated with urbanization

Applying the thermal melanism hypothesis to UHIs predicts pigmentation shifts in urban populations. For example, studies have documented decreased heat-storing pigments in urban wasps [35,37]. However, in *P. rapae*, wing pigmentation interacts with basking behaviour to influence body temperature in complex ways [54–56]. During reflectance basking, highly reflective distal wing regions direct more solar radiation toward the body, such that increased pigmentation in these areas can paradoxically reduce heat gain by limiting reflection. We therefore predicted increased pigmentation at wing tips and wing spots in urban populations, contrary to simple thermal melanism expectations.

However, we found only weak evidence for increased wing tip pigmentation with urbanization and temperature, and no evidence for changes in wing spot pigmentation using an area-based metric. This lack of signal contrasts with experimental evidence showing that pigmentation in these wing areas is plastic in response to temperature and that urban and non-urban populations show evolutionary divergence in pigmentation [38]. Several factors may explain this discrepancy. First, the difference in maximum temperature between urban and non-urban sites in our study (average 2°C based on historical climate data) was smaller than the difference between experimental treatments (26°C vs 29°C) used by [38] and, critically, may not reflect the actual microhabitat temperatures experienced by developing larvae, potentially limiting the expression of plastic responses. Second, wing pigmentation is shaped by multiple selection pressures, including immune stressors [84], and developmental constraints, such as altered larval nutrition [85,86], that may outweigh any thermoregulatory benefits provided by changes in pigmentation. Third, our pigmentation analyses focused on two discrete wing areas involved in thermoregulation, but there may be additional diffuse pigmentation changes across the wing that are difficult to capture using area-based methods, highlighting the need to quantify pigmentation across the entire wing and apply alternative, non-area-based approaches to capture other dimensions of pigmentation variation.

Reflectance properties of butterfly wings vary with climate [31,87] but have not been considered in relation to UHIs. We therefore asked whether visible and NIR reflectance varied with urbanization. Of the seven wing regions analyzed (six NIR, one visible), only the postdiscal forewing showed evidence of urbanization effects, but not as predicted. Instead, we detected the emergence of sexual dimorphism in NIR reflectance in urban populations. Sexual dimorphism in NIR occurs in other butterfly species [31], but our data suggest this dimorphism is absent in non-urban *P. rapae* populations in this wing area. Within each sex, responses to urbanization were weak and in opposing directions. Males showed decreased NIR and visible reflectance with urbanization, which may reduce heat gain when males patrol open areas searching for mates [88], while females showed increased reflectance, the functional significance of which remains unclear and appears maladaptive from a thermal perspective. Although this pattern did not remain significant after FDR correction, it warrants further investigation, particularly because the drivers are unclear. Contrary to our hypothesis that shifts in reflectance track with temperature differences caused by UHIs, our data did not support a role of temperature in driving this sexual dimorphism.

The limited colouration response to urbanization may indicate that *P. rapae* relies primarily on physiological adjustments [89] or behavioural thermoregulation, such as modifying basking postures [55,56] or selecting cooler microhabitats [90], rather than morphological changes to cope with urban heat. Butterflies cease flight when overheating occurs [91], making heat avoidance critical for maintaining foraging and reproductive behaviours, yet such behavioural flexibility may reduce selection for pigmentation or reflectance shifts.

## Conclusions

*P. rapae* shows an affinity for urban habitats that is rare among butterflies, making it an ideal system for studying how urbanization shapes phenotypic evolution. We documented parallel morphological divergence between urban and non-urban populations across three metropolitan regions, with urban butterflies consistently exhibiting reduced body size and altered flight morphology. These results contribute to growing evidence that urbanization drives repeatable change.

While our field observations align closely with plastic responses to temperature documented in experimental studies, we cannot rule out local adaptation, particularly given that *P. rapae* has inhabited North American cities for over 150 years. As urban environments encompass a suite of correlated selection pressures, future work will require integrative approaches, such as common garden experiments and genomic analyses, to disentangle these factors and their evolutionary consequences.

## Supporting information

Electronic supplementary material

## Acknowledgements

Travel for field work was supported by the American Philosophical Society Lewis and Clark Fund and the Entomological Society of America SysEB Student Research Travel Award. CAM was supported by a University Fellowship from Temple University. We thank the Ohio Department of Natural Resources and Pennsylvania Department of Conservation and Natural Resources for access to collection sites. We also thank Tristan Eyring and Henna Desai for assistance with wing size and pigmentation measurements.

