## Supplementary material for "Urbanization drives parallel shifts in morphology and colouration of *Pieris rapae* across three cities": Electronic supplementary material

#### S1. Focal species

*Pieris rapae* is a widespread multivoltine butterfly with established populations on all continents except for South America and Antarctica. Native to Europe, *P. rapae*'s range expanded into Asia in the 1700s before the species was introduced to Africa, North America, and Australia in the 1800s and early 1900s [1]. Historic and genetic data support at least one introduction from Europe to North America around 1855 [1,2]. Therefore, this species has been in North America since the Industrial Revolution and growth of cities. A habitat generalist, *P. rapae* can be found in multiple environments, from forest edges and agricultural landscapes to disturbed urban areas [3,4]. Caterpillars feed exclusively on plants in the Brassicaceae family [5,6], and, like many other butterflies, adult *P. rapae* are nectar generalists [5]. Female *P. rapae* can be distinguished from males based on the presence of an additional eyespot and darker wing pigmentation [7].

Although *P. rapae* is nonmigratory across much of its range, including North America, it is nonetheless considered highly mobile [8]. However, movement behaviour differs between sexes, suggesting they could face different challenges in urban environments. Newly emerged females mate at their natal sites [9,10] and subsequently disperse widely as they lay eggs across multiple host plant patches [11,12]. Studies of Australian and Japanese populations found female *P. rapae* moved a net distance of 2 km on average over their lifetime [10,13]. In contrast, males remain near their emergence sites, with extensive daily, local movements associated with mate-patrolling behaviour [14].

#### S2. Study site selection and characterization

##### S2.1 City descriptions

Philadelphia is the oldest and largest of the three cities, founded in 1682, and covers 347 $km^2$ with a current city population of over 1.6 million at a density of ~4,600 people/$km^2$ [15]. There are multiple smaller disjunct urban areas within the metropolitan region, one of which is Conshohocken, which we included as an additional urban site within Philadelphia. Pittsburgh was founded in 1758 and covers ~150 $km^2$ with a population close to 300,000 and density of 2,000 people/$km^2$ [15]. Cleveland was most recently founded in 1796, covers ~200 $km^2$, and has a population of approximately 370,000 and density of 3,800 people/$km^2$ [15]. The cities also differ in climate despite similar latitudes (Cleveland: 41.4°N, Pittsburgh and Philadelphia: 40.4°N). Philadelphia is considerably warmer than Pittsburgh and Cleveland and is classified as a moist subtropical climate, whereas Pittsburgh and Cleveland share a moist continental climate [16].

**S2.2 Site selection criteria**

In each region, we identified three urban sites (with one additional urban site in Philadelphia) and three non-urban sites (additional site information in electronic supplementary material, S2.2 and table S1). Sites were 0.75 km radius circles centered on a single GPS coordinate (pairwise site distances in electronic supplementary material, S2.2 and table S1). Urban sites were defined as those with at least 50% impervious surface cover (following [17]), based on the National Land Cover Database (NLCD) Fractional Impervious Surface data [18]. Most urban sites were residential areas, including small community gardens, vacant lots, and lawns, whereas non-urban sites were primarily within state or county parks and consisted of forest edges and grasslands.

Urban sites within the same region were separated by at least 7 km (table S1). This distance balanced the logistical challenges of finding multiple suitable urban sites sufficiently spaced apart, which can be difficult depending on city size and layout, while still allowing us to sample replicate sites within the same metropolitan area. Non-urban sites surrounded each city and were at least 20 km from urban sites and from each other. Pairwise distances between the centers of sites were calculated as Haversine distances using the R package geosphere (v. 1.5.20) [19].

**Table S1.** Pairwise distances (km) among sampling sites.

| | CLEV-NU1 | CLEV-NU2 | CLEV-NU3 | CLEV-U1 | CLEV-U2 | CLEV-U3 | PITT-NU1 | PITT-NU2 | PITT-NU3 | PITT-U1 | PITT-U2 | PITT-U3 | PHIL-NU1 | PHIL-NU2 | PHIL-NU3 | PHIL-U1 | PHIL-U2 | PHIL-U3 |
|---|---|---|---|---|---|---|---|---|---|---|---|---|---|---|---|---|---|---|
| **CLEV-NU2** | 100.9 | | | | | | | | | | | | | | | | | |
| **CLEV-NU3** | 89.5 | 21.8 | | | | | | | | | | | | | | | | |
| **CLEV-U1** | 67.5 | 51.7 | 56.2 | | | | | | | | | | | | | | | |
| **CLEV-U2** | 52.3 | 58.7 | 57.5 | 15.3 | | | | | | | | | | | | | | |
| **CLEV-U3** | 58.0 | 59.3 | 60.8 | 10.2 | 7.5 | | | | | | | | | | | | | |
| **PITT-NU1** | 168.4 | 106.6 | 95.9 | 152.2 | 152.0 | 156.3 | | | | | | | | | | | | |
| **PITT-NU2** | 179.6 | 90.3 | 91.7 | 142.0 | 147.4 | 149.1 | 55.4 | | | | | | | | | | | |
| **PITT-NU3** | 253.0 | 174.9 | 171.8 | 225.8 | 229.2 | 232.0 | 86.9 | 86.7 | | | | | | | | | | |
| **PITT-U1** | 206.8 | 134.3 | 128.3 | 183.7 | 185.7 | 189.1 | 40.3 | 55.0 | 46.7 | | | | | | | | | |
| **PITT-U2** | 213.9 | 142.5 | 136.3 | 191.8 | 193.6 | 197.1 | 46.3 | 62.8 | 40.9 | 8.3 | | | | | | | | |
| **PITT-U3** | 199.1 | 129.0 | 122.0 | 177.7 | 179.2 | 182.8 | 31.8 | 54.2 | 55.1 | 8.7 | 14.8 | | | | | | | |
| **PHIL-NU1** | 587.0 | 496.2 | 500.0 | 547.3 | 554.7 | 555.5 | 424.8 | 408.3 | 338.0 | 384.5 | 378.8 | 393.0 | | | | | | |
| **PHIL-NU2** | 582.4 | 489.8 | 494.6 | 540.6 | 548.5 | 549.0 | 422.6 | 403.1 | 336.2 | 382.3 | 377.1 | 390.9 | 24.2 | | | | | |
| **PHIL-NU3** | 599.5 | 505.4 | 511.1 | 555.6 | 564.0 | 564.2 | 442.1 | 419.9 | 356.2 | 401.9 | 397.0 | 410.5 | 47.9 | 27.8 | | | | |
| **PHIL-U1** | 609.9 | 518.5 | 522.6 | 569.4 | 577.0 | 577.7 | 448.4 | 430.9 | 361.7 | 408.1 | 402.5 | 416.6 | 24.4 | 31.8 | 40.0 | | | |
| **PHIL-U2** | 615.6 | 523.3 | 527.9 | 574.0 | 581.9 | 582.4 | 455.2 | 436.3 | 368.6 | 414.9 | 409.5 | 423.5 | 35.1 | 33.5 | 31.3 | 13.3 | | |
| **PHIL-U3** | 609.5 | 517.4 | 521.9 | 568.2 | 576.0 | 576.6 | 448.8 | 430.3 | 362.1 | 408.5 | 403.0 | 417.0 | 28.1 | 28.1 | 31.7 | 8.6 | 7.0 | |
| **PHIL-U4** | 595.8 | 503.7 | 508.2 | 554.5 | 562.3 | 562.9 | 435.2 | 416.5 | 348.6 | 394.9 | 389.5 | 403.4 | 19.2 | 15.1 | 29.5 | 16.7 | 20.1 | 13.7 |

**S2.3 Environmental data**

Site data were calculated within concentric circular buffers (0.75, 1, 1.5, 2, and 2.5 km radii) centered on the site centroid. The 750 m buffer represents the size of the sites themselves. We excluded buffers >2.5 km to avoid overlap among urban sites and to prevent issues with obtaining environmental data for sites close to bodies of water (CLEV-U1 and PHIL-U1).

We quantified impervious surface within buffers using the National Land Cover Database (NLCD) Fractional Impervious Surface product, which reports the proportion of impervious surface cover within each 30 m pixel [18]. We also calculated habitat heterogeneity within each buffer by estimating perimeter area ratio (PARA) using the R package landscapemetrics [20]. To calculate PARA, we used the impervious surface dataset to categorize each pixel as developed (>20% impervious surface) or undeveloped (≤20% impervious surface), based on cutoffs used in NLCD's Land Cover classifications [18]. PARA calculations are based only on the undeveloped patches, following [21]. High PARA values indicate highly subdivided, edge-dominated remaining undeveloped areas, while low values indicate larger, more intact blocks of undeveloped areas. PARA could not be calculated for one urban site (PHIL-U1) for 750 m and 1 km buffers because there was no undeveloped land cover, resulting in a denominator of zero.

Lastly, we obtained mean and maximum near-surface air temperature measurements from the Global Seamless High-resolution Temperature Dataset [22] for June, July, and August of each year, as this coincides with our collections and the timing of the summer form of *P. rapae*. This product provides daily temperature measurements at 30 arcsecond resolution.

We calculated site-level means for impervious surface, PARA, and temperature based on temporal averages (2000-2022 for impervious surface and PARA, 2001-2020 for temperature) using R packages terra (v. 1.8.29) [23] and sf (v. 1.0.19) [24,25].

**S2.4 Buffer size selection**

We calculated pairwise Pearson correlation coefficients ($r$) among buffer sizes to determine whether nested buffers provided distinct environmental information. The nested buffers around sites were highly correlated for each environmental variable ($r$ = 0.76–1.0, table S2). We elected to use a 1 km radius buffer considering the typical lifetime movement of *P. rapae* and the fact that urbanization has been found to drive phenotypic change in butterflies at scales equal to or below a species' dispersal distance [26].

**Table S2.** Correlations across circular buffer sizes (varying radii) for each environmental variable.

**Percent Impervious Surface Cover**

| | 750 m | 1 km | 1.5 km | 2 km |
|---|---|---|---|---|
| 1 km | 1.00 | -- | -- | -- |
| 1.5 km | 0.98 | 0.99 | -- | -- |
| 2 km | 0.97 | 0.98 | 1.00 | -- |
| 2.5 km | 0.96 | 0.98 | 0.99 | 1.00 |

**Mean Summer Temperature (°C)**

| | 750 m | 1 km | 1.5 km | 2 km |
|---|---|---|---|---|
| 1 km | 1.00 | -- | -- | -- |
| 1.5 km | 1.00 | 1.00 | -- | -- |
| 2 km | 1.00 | 1.00 | 1.00 | -- |
| 2.5 km | 1.00 | 1.00 | 1.00 | 1.00 |

**Max. Summer Temperature (°C)**

| | 750 m | 1 km | 1.5 km | 2 km |
|---|---|---|---|---|
| 1 km | 1.00 | -- | -- | -- |
| 1.5 km | 1.00 | 1.00 | -- | -- |
| 2 km | 1.00 | 1.00 | 1.00 | -- |
| 2.5 km | 1.00 | 1.00 | 1.00 | 1.00 |

**Perimeter-Area Ratio**

| | 750 m | 1 km | 1.5 km | 2 km |
|---|---|---|---|---|
| 1 km | 0.77 | -- | -- | -- |
| 1.5 km | 0.78 | 0.90 | -- | -- |
| 2 km | 0.80 | 0.83 | 0.96 | -- |
| 2.5 km | 0.76 | 0.84 | 0.93 | 0.98 |

**S2.5 Environmental variable reduction**

We did not proceed with impervious surface as a continuous proxy for urbanization as its distribution was strongly bimodal and exhibited minimal variation within each site type. We examined correlations among environmental variables to identify highly correlated metrics for exclusion. Among site variables, mean and maximum air temperature were highly correlated within regions ($r$ = 0.92–1.0, table S3). We used maximum air temperature (hereafter, “temperature”) for our analyses as it has been shown to more strongly impact fitness of another Pieridae species than mean temperature [27]. Temperature and PARA were also highly correlated across sites within regions ($r$ =

0.86–0.95, table S3), indicating that both variables captured the same underlying environmental gradient. We retained temperature as the representative variable for this shared gradient.

**Table S3.** Correlations among environmental variables within each region.

**Cleveland**

| | Impervious Surface | Mean Temperature | Max. Temperature |
|---|---|---|---|
| Mean Temperature | 0.98 | -- | -- |
| Max. Temperature | 0.86 | 0.92 | -- |
| Perimeter-Area Ratio | 0.86 | 0.89 | 0.92 |

**Pittsburgh**

| | Impervious Surface | Mean Temperature | Max. Temperature |
|---|---|---|---|
| Mean Temperature | 1.00 | -- | -- |
| Max. Temperature | 1.00 | 1.00 | -- |
| Perimeter-Area Ratio | 0.85 | 0.84 | 0.86 |

**Philadelphia**

| | Impervious Surface | Mean Temperature | Max. Temperature |
|---|---|---|---|
| Mean Temperature | 0.99 | -- | -- |
| Max. Temperature | 0.95 | 0.98 | -- |
| Perimeter-Area Ratio | 0.90 | 0.92 | 0.95 |

**Table S4.** Sampling site information. Environmental data were calculated within 1 km buffers. % Imp. = percent impervious surface cover; Tmean, Tmax = mean and maximum summer temperatures (June–August); PARA = perimeter-area ratio of undeveloped patches.

| Site Code | Site Name | Region | Site Type | Lat. | Long. | % Imp. | Tmean | Tmax | PARA | Males | Females |
|---|---|---|---|---|---|---|---|---|---|---|---|
| CLEV-NU1 | Findley | Cleveland | Non-urban | 41.1323 | -82.2145 | 1.8 | 21.5 | 28.5 | 24.6 | 20 | 10 |
| CLEV-NU2 | Nelson Kennedy | Cleveland | Non-urban | 41.3286 | -81.0383 | 2.3 | 21.0 | 28.3 | 13.7 | 22 | 9 |
| CLEV-NU3 | West Branch | Cleveland | Non-urban | 41.1506 | -81.1477 | 5.1 | 21.1 | 28.2 | 45.6 | 12 | 9 |
| CLEV-U1 | University Circle | Cleveland | Urban | 41.5248 | -81.5994 | 63.5 | 23.1 | 30.0 | 547.5 | 22 | 2 |
| CLEV-U2 | Brooklyn | Cleveland | Urban | 41.4272 | -81.7281 | 62.3 | 23.1 | 30.6 | 402.8 | 23 | 4 |
| CLEV-U3 | Downtown | Cleveland | Urban | 41.4936 | -81.7140 | 59.6 | 22.7 | 29.0 | 165.7 | 8 | 4 |
| PITT-NU1 | Raccoon Creek | Pittsburgh | Non-urban | 40.5024 | -80.3972 | 1.0 | 21.3 | 28.3 | 7.7 | 20 | 14 |
| PITT-NU2 | Moraine | Pittsburgh | Non-urban | 40.9428 | -80.0904 | 2.6 | 21.2 | 28.1 | 27.3 | 21 | 7 |
| PITT-NU3 | Keystone | Pittsburgh | Non-urban | 40.3771 | -79.3847 | 3.5 | 21.5 | 28.5 | 21.6 | 26 | 16 |
| PITT-U1 | Shadyside | Pittsburgh | Urban | 40.4653 | -79.9235 | 64.7 | 23.5 | 31.1 | 487.1 | 21 | 15 |
| PITT-U2 | Braddock | Pittsburgh | Urban | 40.4050 | -79.8657 | 53.3 | 23.2 | 30.6 | 144.1 | 20 | 16 |
| PITT-U3 | Manchester | Pittsburgh | Urban | 40.4583 | -80.0256 | 62.8 | 23.5 | 30.9 | 237.9 | 4 | 7 |
| PHIL-NU1 | Ridley | Philadelphia | Non-urban | 39.9467 | -75.4518 | 1.3 | 22.9 | 30.2 | 3.4 | 27 | 8 |
| PHIL-NU2 | Evansburg | Philadelphia | Non-urban | 40.1641 | -75.4359 | 4.4 | 23.2 | 30.6 | 24.2 | 29 | 4 |
| PHIL-NU3 | Peace Valley | Philadelphia | Non-urban | 40.3255 | -75.1858 | 2.7 | 22.9 | 30.0 | 6.6 | 28 | 7 |
| PHIL-U1 | North | Philadelphia | Urban | 39.9664 | -75.1667 | 75.9 | 24.9 | 32.0 | NA | 25 | 4 |
| PHIL-U2 | Northeast | Philadelphia | Urban | 40.0592 | -75.0676 | 60.3 | 24.5 | 32.2 | 394.1 | 33 | 5 |
| PHIL-U3 | Northwest | Philadelphia | Urban | 40.0421 | -75.1471 | 64.5 | 24.8 | 32.3 | 332.3 | 24 | 11 |
| PHIL-U4 | Conshohocken | Philadelphia | Urban | 40.0754 | -75.3021 | 55.8 | 24.3 | 31.5 | 133.9 | 24 | 10 |

### S3. Protocols for morphological measurements

#### S3.1 Photography setup

Wings were photographed with a Nikon D3300 with AF-S DX NIKKOR 18-55mm f/3.5-5.6G VR II lens with the aperture set to f/13. The camera was at a constant height with a level to ensure the camera stayed horizontal. Photos were illuminated with artificial light, and there was no external natural light. We placed a label with the sample ID, a ruler, and a colour reference card within the frame of each photograph. Wings were photographed on an 18% grey standard.

#### S3.2 Wing reflectance

Wing reflectance was captured using a UV-NIR spectrometer (Ocean Optics; Orlando FL, USA) equipped with a deuterium-tungsten light source. Preliminary tests holding the probe at 45° vs 90° revealed no difference in reflectance. All reflectance measurements were captured with the probe at 90° (perpendicular) to the wing, as this provided more precise positioning. Figure S1 shows areas on the wings where reflectance was measured.

Spectra were processed using the R package pavo (v. 2.9.0) [28]. We applied LOESS smoothing to reduce spectral noise and a negative value correction that converted negative values to zero. The average percent reflectance ("B2" from pavo's summary function) was calculated separately over the ultraviolet (UV, 250-400 nm), visible (400-700 nm), and near infrared (NIR, 700-950 nm) portions of the spectrum for each wing spot. Since UV wavelengths are relatively unimportant for temperature regulation and more vital for sexual signaling in butterflies [29], we omitted UV measurements from our analyses and only report results for NIR and visible reflectance (hereafter, NIR-r and Vis-r).

We considered the use of principal component analysis to reduce reflectance measurements to composite metrics. However, because males and females exhibited contrasting patterns of reflectance, the resulting components were not interpretable; therefore, we do not present these analyses.

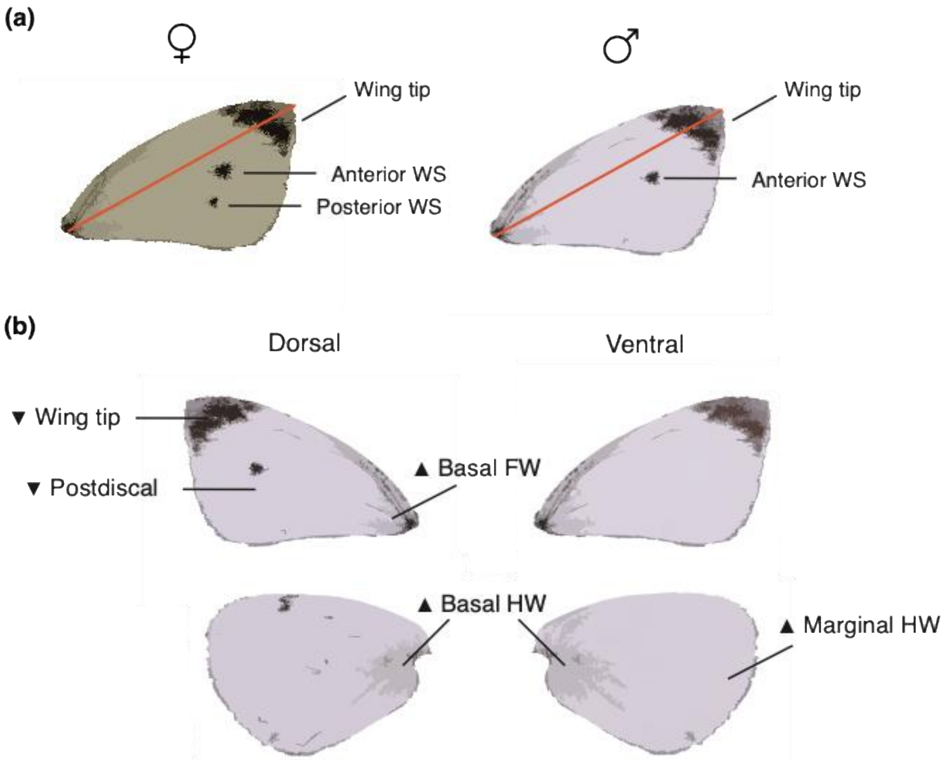

**Figure S1.** *Pieris rapae* wings showing measurement locations for size, pigmentation, and reflectance traits. (a) Dorsal forewings of female (left) and male (right), showing forewing length (red line) and pigmented areas measured (WS = wing spot). All pigmented areas were predicted to increase in urban populations. (b) Dorsal (left) and ventral (right) fore- and hindwings (FW, HW). Near-infrared and visible reflectance were measured for labeled areas via spectrometry. ▲ = reflectance predicted to increase in urban butterflies; ▼ = reflectance predicted to decrease in urban butterflies.

#### S4. Statistical analyses

##### S4.1 Variable reduction

Among morphological traits, total mass and thorax mass were strongly correlated within regions ($r$ = 0.71–0.81, figure S2a). However, we retained both total mass as our primary measure of overall body size and thorax mass because it reflects investment in flight musculature. We distinguished reductions in thorax size from overall body size reductions by accounting for total mass in thorax mass analyses. Wing loading was likewise strongly correlated with total mass and was excluded ($r$ = 0.89–0.90, figure S2a). To isolate potential effects on aerodynamics that are independent of overall body size, we retained forewing area (the numerator of wing loading) and adjusted for total mass in analyses. As wing length and area were highly correlated ($r$ = 0.91–0.93, figure S2a), we also excluded wing length.

Area of the anterior and posterior wing spots were strongly correlated within regions ($r$ = 0.75–0.86, figure S2b). Because the posterior spot is absent in males, we retained only the anterior spot. VIS-r and NIR-r were also highly correlated in all wing areas except for the postdiscal area (figure S2c). We therefore analyzed both Vis-r and NIR-r in the postdiscal region, but restricted analyses to NIR-r in all other areas given its stronger relevance to thermoregulation [30].

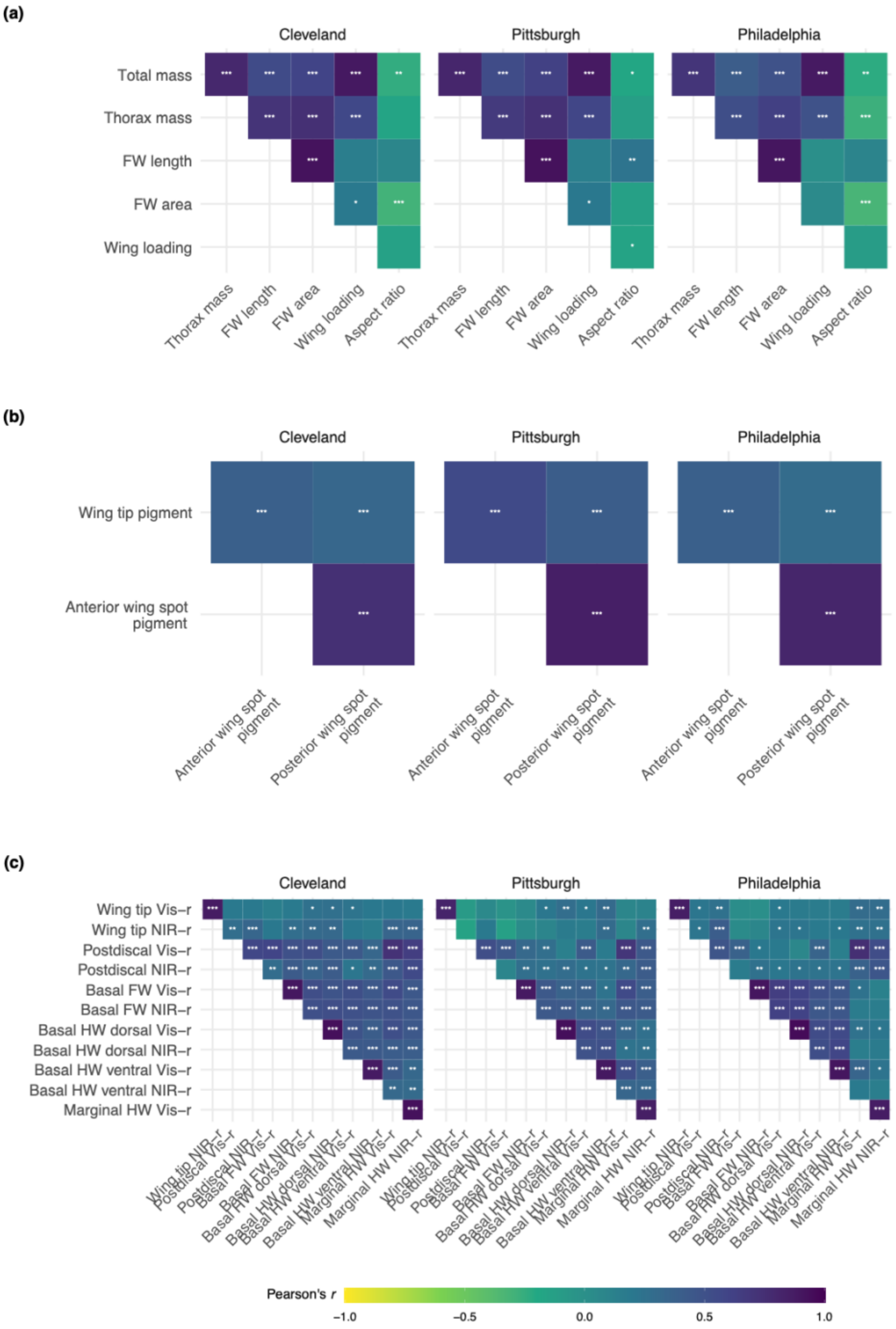

**Figure S2.** Correlations among morphological traits in *Pieris rapae*, with significance indicated (* = p < 0.05, ** = p < 0.01, *** = p < 0.001). Traits are grouped into trait categories: (a) size and mobility, (b) pigmentation, and (b) NIR reflectance (NIR-r) and visible reflectance (Vis-r).

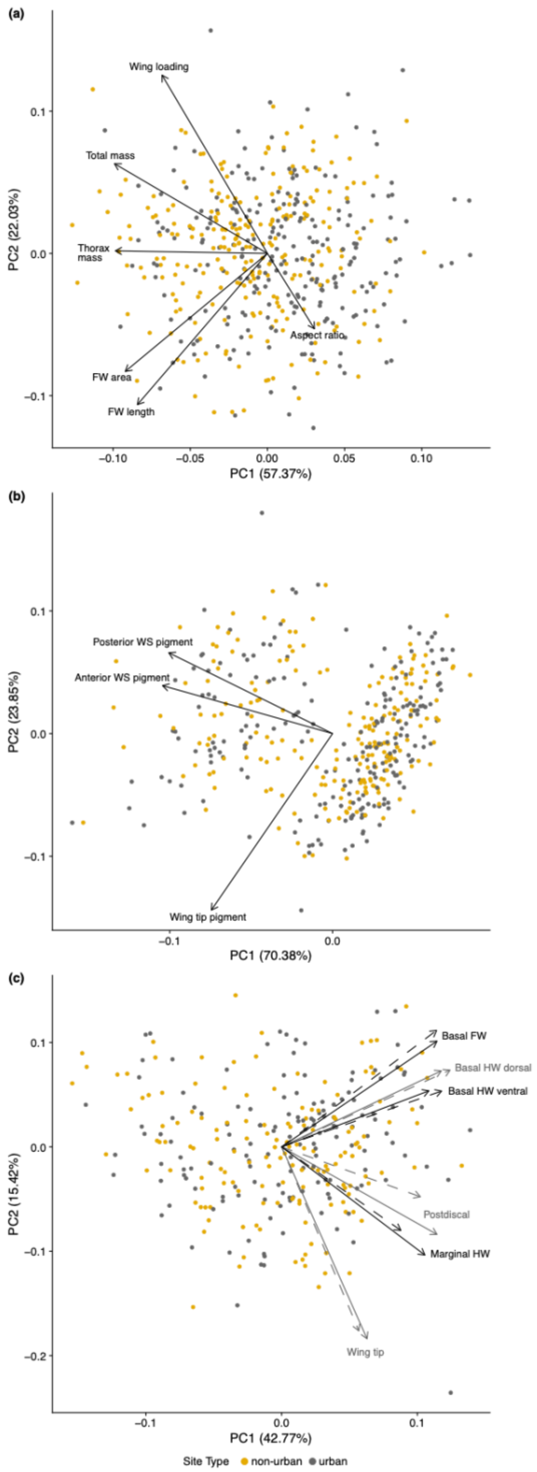

**Figure S3.** Trait PCA biplots showing (a) morphometrics, (b) wing pigmentation, and (c) wing reflectance. Each point represents an individual butterfly (gold = non-urban, grey = urban). Arrows indicate trait loadings on PC1 and PC2. For reflectance traits, each wing region has two loadings: dashed lines for visible reflectance (400-700 nm) and solid lines for near-infrared reflectance (700-950 nm). Abbreviations: WS = wing spot, FW = forewing, HW = hindwing.

**Table S5.** Sample sizes per trait measurement.

**Size**

| | **Total mass** | | **Thorax mass** | | **Wing area** | | **Aspect ratio** | |
|---|---|---|---|---|---|---|---|---|
| Site | M | F | M | F | M | F | M | F |
| CLEV-NU1 | 20 | 10 | 20 | 10 | 20 | 7 | 20 | 7 |
| CLEV-NU2 | 22 | 8 | 22 | 8 | 19 | 7 | 19 | 7 |
| CLEV-NU3 | 12 | 9 | 12 | 9 | 11 | 6 | 11 | 6 |
| CLEV-U1 | 22 | 2 | 22 | 2 | 19 | 1 | 19 | 1 |
| CLEV-U2 | 23 | 4 | 23 | 4 | 23 | 3 | 23 | 3 |
| CLEV-U3 | 8 | 4 | 8 | 4 | 7 | 4 | 7 | 4 |
| PITT-NU1 | 20 | 14 | 20 | 14 | 16 | 12 | 16 | 12 |
| PITT-NU2 | 20 | 7 | 20 | 7 | 19 | 5 | 20 | 5 |
| PITT-NU3 | 26 | 13 | 26 | 13 | 23 | 11 | 23 | 14 |
| PITT-U1 | 21 | 15 | 21 | 15 | 21 | 12 | 21 | 12 |
| PITT-U2 | 20 | 16 | 20 | 16 | 17 | 12 | 17 | 12 |
| PITT-U3 | 4 | 6 | 4 | 6 | 3 | 6 | 3 | 7 |
| PHIL-NU1 | 26 | 7 | 26 | 7 | 25 | 4 | 25 | 4 |
| PHIL-NU2 | 29 | 4 | 29 | 4 | 24 | 2 | 24 | 2 |
| PHIL-NU3 | 28 | 7 | 28 | 7 | 27 | 6 | 27 | 6 |
| PHIL-U1 | 24 | 4 | 24 | 4 | 23 | 4 | 23 | 4 |
| PHIL-U2 | 31 | 5 | 31 | 5 | 25 | 5 | 26 | 5 |
| PHIL-U3 | 24 | 10 | 24 | 10 | 21 | 8 | 21 | 8 |
| PHIL-U4 | 24 | 10 | 24 | 10 | 20 | 9 | 20 | 9 |

**Pigmentation**

| Site | Wing tip pigment area | | Wing spot pigment area | |
|---|---|---|---|---|
| | M | F | M | F |
| CLEV-NU1 | 18 | 7 | 19 | 7 |
| CLEV-NU2 | 12 | 6 | 14 | 7 |
| CLEV-NU3 | 9 | 6 | 11 | 6 |
| CLEV-U1 | 16 | 1 | 16 | 1 |
| CLEV-U2 | 16 | 3 | 18 | 3 |
| CLEV-U3 | 5 | 3 | 5 | 4 |
| PITT-NU1 | 13 | 11 | 14 | 12 |
| PITT-NU2 | 18 | 5 | 20 | 5 |
| PITT-NU3 | 17 | 12 | 19 | 13 |
| PITT-U1 | 17 | 12 | 16 | 12 |
| PITT-U2 | 11 | 11 | 12 | 12 |
| PITT-U3 | 1 | 7 | 1 | 7 |
| PHIL-NU1 | 20 | 2 | 20 | 4 |
| PHIL-NU2 | 12 | 2 | 13 | 2 |
| PHIL-NU3 | 14 | 5 | 18 | 5 |
| PHIL-U1 | 14 | 4 | 15 | 4 |
| PHIL-U2 | 16 | 5 | 17 | 5 |
| PHIL-U3 | 17 | 8 | 17 | 8 |
| PHIL-U4 | 14 | 9 | 15 | 9 |

**Reflectance**

| Site | Wing tip NIR-r | | Post-discal NIR-r | | Post-discal Vis-r | | Basal FW NIR-r | | Basal HW dorsal NIR-r | | Basal HW ventral NIR-r | | Marginal HW NIR-r | |
|---|---|---|---|---|---|---|---|---|---|---|---|---|---|---|
| | M | F | M | F | M | F | M | F | M | F | M | F | M | F |
| CLEV-NU1 | 10 | 8 | 10 | 10 | 10 | 10 | 10 | 10 | 11 | 10 | 11 | 10 | 11 | 10 |
| CLEV-NU2 | 10 | 9 | 10 | 9 | 10 | 9 | 10 | 9 | 10 | 9 | 10 | 9 | 10 | 9 |
| CLEV-NU3 | 10 | 9 | 10 | 9 | 10 | 9 | 10 | 9 | 10 | 9 | 10 | 9 | 10 | 8 |
| CLEV-U1 | 9 | 2 | 9 | 2 | 9 | 2 | 9 | 2 | 9 | 2 | 9 | 2 | 9 | 2 |
| CLEV-U2 | 10 | 4 | 10 | 4 | 10 | 4 | 10 | 4 | 10 | 4 | 10 | 4 | 9 | 4 |
| CLEV-U3 | 6 | 4 | 6 | 4 | 6 | 4 | 6 | 4 | 6 | 4 | 6 | 4 | 6 | 4 |
| PITT-NU1 | 9 | 10 | 9 | 9 | 9 | 9 | 9 | 10 | 9 | 9 | 9 | 10 | 9 | 10 |
| PITT-NU2 | 10 | 6 | 10 | 6 | 10 | 6 | 10 | 6 | 10 | 6 | 10 | 6 | 10 | 6 |
| PITT-NU3 | 9 | 10 | 9 | 10 | 9 | 10 | 9 | 10 | 10 | 10 | 10 | 10 | 10 | 10 |
| PITT-U1 | 10 | 10 | 10 | 10 | 10 | 10 | 10 | 10 | 9 | 10 | 9 | 10 | 9 | 9 |
| PITT-U2 | 10 | 10 | 10 | 10 | 10 | 10 | 10 | 10 | 10 | 10 | 10 | 10 | 10 | 10 |
| PITT-U3 | 2 | 7 | 2 | 7 | 2 | 7 | 2 | 7 | 2 | 7 | 2 | 7 | 2 | 7 |
| PHIL-NU1 | 10 | 8 | 10 | 8 | 10 | 8 | 10 | 8 | 10 | 8 | 10 | 8 | 10 | 7 |
| PHIL-NU2 | 10 | 4 | 10 | 4 | 10 | 4 | 10 | 4 | 10 | 4 | 10 | 4 | 10 | 3 |
| PHIL-NU3 | 8 | 5 | 8 | 5 | 8 | 5 | 8 | 5 | 7 | 5 | 7 | 5 | 7 | 5 |
| PHIL-U1 | 10 | 4 | 10 | 4 | 10 | 4 | 10 | 4 | 10 | 4 | 10 | 4 | 10 | 4 |
| PHIL-U2 | 9 | 5 | 9 | 5 | 9 | 5 | 9 | 5 | 9 | 5 | 9 | 5 | 9 | 5 |
| PHIL-U3 | 7 | 10 | 7 | 10 | 7 | 10 | 7 | 10 | 7 | 10 | 7 | 10 | 7 | 10 |
| PHIL-U4 | 10 | 10 | 10 | 10 | 10 | 10 | 10 | 10 | 10 | 10 | 10 | 10 | 10 | 10 |

##### S4.2 Testing the impacts of urbanization

We initially fit models with lme4 (v. 1.1.36) [31] and evaluated model assumptions using DHARMa (v. 0.4.7) [32]. Because we detected heteroscedasticity, we refit models using nlme (v. 3.1.166) [33,34] with a weighting function that allowed residual variances to differ across levels of the grouping factor exhibiting heteroscedasticity. For all models, we performed joint tests for fixed effects using emmeans (v. 1.10.6) [35].

#### S5. Supplemental results

Tables S6-S8 provide model coefficients and test statistics for morphometric, pigmentation, and reflectance analyses. Both raw and false discovery rate (FDR) corrected *p* values are reported.

**Table S6.** Model coefficients for size traits. UvNU = urban vs non-urban; ΔAIC = temperature model AIC minus urban model AIC; FDR-corrected p-values in parentheses.

| Trait | Model | Effect | numDF | denDF | *F* | *p* |
|---|---|---|---|---|---|---|
| Total mass | Urban vs non-urban | Urban vs non-urban | 1 | 13 | 16.00 | 0.002 (0.006) |
| | | Region | 2 | 13 | 11.93 | 0.001 (0.002) |
| | | Sex | 1 | 538 | 0.001 | 0.98 (0.98) |
| | | UvNU:Region | 2 | 13 | 1.36 | 0.29 (0.55) |
| | | UvNU:Sex | 1 | 538 | 3.33 | 0.07 (0.15) |
| | Temperature gradient | Temperature | 1 | 13 | 21.53 | 0.001 (0.002) |
| | | Region | 2 | 13 | 20.00 | < 0.001 (< 0.001) |
| | | Sex | 1 | 538 | 0.00 | 0.96 (0.96) |
| | | Temperature:Region | 2 | 13 | 0.37 | 0.70 (0.70) |
| | | Temperature:Sex | 1 | 538 | 2.49 | 0.12 (0.23) |
| Model comparison: Urban vs non-urban AIC = 3132.8, Temperature AIC = 3138.4, ΔAIC = 5.6 | | | | | | |
| Thorax mass | Urban vs non-urban | Urban vs non-urban | 1 | 13 | 5.46 | 0.04 (0.07) |
| | | Region | 2 | 13 | 2.85 | 0.09 (0.09) |
| | | Sex | 1 | 537 | 229.72 | < 0.001 (< 0.001) |
| | | Total mass | 1 | 537 | 1,039.93 | < 0.001 |
| | | UvNU:Region | 2 | 13 | 2.39 | 0.13 (0.52) |
| | | UvNU:Sex | 1 | 537 | 0.08 | 0.78 (0.78) |
| | Temperature gradient | Temperature | 1 | 13 | 7.60 | 0.02 (0.03) |
| | | Region | 2 | 13 | 2.36 | 0.13 (0.13) |
| | | Sex | 1 | 537 | 225.68 | < 0.001 (< 0.001) |
| | | Total mass | 1 | 537 | 1,038.73 | < 0.001 |
| | | Temperature:Region | 2 | 13 | 3.11 | 0.08 (0.31) |
| | | Temperature:Sex | 1 | 537 | 0.47 | 0.49 (0.55) |
| Model comparison: Urban vs non-urban AIC = 1349.7, Temperature AIC = 1357.5, ΔAIC = 7.8 | | | | | | |

| Trait | Model | Effect | numDF | denDF | *F* | *p* |
| --- | --- | --- | --- | --- | --- | --- |
| Wing area | Urban vs non-urban | Urban vs non-urban | 1 | 13 | 0.51 | 0.49 (0.65) |
| | | Region | 2 | 13 | 18.63 | < 0.001 (0.001) |
| | | Sex | 1 | 465 | 89.63 | < 0.001 (< 0.001) |
| | | Total mass | 1 | 465 | 244.61 | < 0.001 |
| | | UvNU:Region | 2 | 13 | 0.64 | 0.54 (0.55) |
| | | UvNU:Sex | 1 | 465 | 3.23 | 0.07 (0.15) |
| | Temperature gradient | Temperature | 1 | 13 | 2.11 | 0.17 (0.23) |
| | | Region | 2 | 13 | 16.67 | < 0.001 (0.001) |
| | | Sex | 1 | 465 | 80.76 | < 0.001 (< 0.001) |
| | | Total mass | 1 | 465 | 242.17 | < 0.001 |
| | | Temperature:Region | 2 | 13 | 1.39 | 0.28 (0.57) |
| | | Temperature:Sex | 1 | 465 | 3.93 | 0.05 (0.19) |
| Model comparison: Urban vs non-urban AIC = 4084.4, Temperature AIC = 4089.1, ΔAIC = 4.7 | | | | | | |
| Aspect ratio | Urban vs non-urban | Urban vs non-urban | 1 | 13 | 0.001 | 0.97 (0.97) |
| | | Region | 2 | 13 | 3.78 | 0.05 (0.07) |
| | | Sex | 1 | 472 | 8.85 | 0.003 (0.004) |
| | | UvNU:Region | 2 | 13 | 0.62 | 0.55 (0.55) |
| | | UvNU:Sex | 1 | 472 | 0.72 | 0.40 (0.53) |
| | Temperature gradient | Temperature | 1 | 13 | 0.15 | 0.70 (0.70) |
| | | Region | 2 | 13 | 2.88 | 0.09 (0.12) |
| | | Sex | 1 | 472 | 8.98 | 0.003 (0.004) |
| | | Temperature:Region | 2 | 13 | 0.67 | 0.53 (0.70) |
| | | Temperature:Sex | 1 | 472 | 0.35 | 0.55 (0.55) |
| Model comparison: Urban vs non-urban AIC = 618.9, Temperature AIC = 625.7, ΔAIC = 6.8 | | | | | | |

**Table S7.** Model coefficients for pigmentation traits. UvNU = urban vs non-urban; ΔAIC = temperature model AIC minus urban model AIC; FDR-corrected p-values in parentheses.

| Trait | Model | Effect | numDF | denDF | *F* | *p* |
|---|---|---|---|---|---|---|
| Wing tip pigment area | Urban vs non-urban | Urban vs non-urban | 1 | 13 | 3.57 | 0.08 (0.16) |
| | | Region | 2 | 13 | 13.90 | 0.001 (0.001) |
| | | Sex | 1 | 357 | 75.10 | < 0.001 (< 0.001) |
| | | Forewing area | 1 | 357 | 27.19 | < 0.001 |
| | | UvNU:Region | 2 | 13 | 1.02 | 0.39 (0.55) |
| | | UvNU:Sex | 1 | 357 | 0.18 | 0.68 (0.68) |
| | Temperature gradient | Temperature | 1 | 13 | 3.12 | 0.10 (0.20) |
| | | Region | 2 | 13 | 10.28 | 0.002 (0.004) |
| | | Sex | 1 | 357 | 75.79 | < 0.001 (< 0.001) |
| | | Forewing area | 1 | 357 | 27.24 | < 0.001 |
| | | Temperature:Region | 2 | 13 | 1.14 | 0.35 (0.36) |
| | | Temperature:Sex | 1 | 357 | 0.01 | 0.92 (0.92) |
| Model comparison: Urban vs non-urban AIC = 2095.2, Temperature AIC = 2102.8, ΔAIC = 7.6 | | | | | | |
| Wing spot pigment area | Urban vs non-urban | Urban vs non-urban | 1 | 13 | 0.32 | 0.58 (0.58) |
| | | Region | 2 | 13 | 0.68 | 0.53 (0.53) |
| | | Sex | 1 | 384 | 670.77 | < 0.001 (< 0.001) |
| | | Forewing area | 1 | 384 | 40.68 | < 0.001 |
| | | UvNU:Region | 2 | 13 | 0.62 | 0.55 (0.55) |
| | | UvNU:Sex | 1 | 384 | 0.42 | 0.52 (0.68) |
| | Temperature gradient | Temperature | 1 | 13 | 0.35 | 0.56 (0.56) |
| | | Region | 2 | 13 | 0.95 | 0.41 (0.41) |
| | | Sex | 1 | 384 | 671.00 | < 0.001 (< 0.001) |
| | | Forewing area | 1 | 384 | 38.95 | < 0.001 |
| | | Temperature:Region | 2 | 13 | 1.10 | 0.36 (0.36) |
| | | Temperature:Sex | 1 | 384 | 2.75 | 0.10 (0.20) |
| Model comparison: Urban vs non-urban AIC = 829.2, Temperature AIC = 832.5, ΔAIC = 3.3 | | | | | | |

**Table S8.** Model coefficients for reflectance traits. UvNU = urban vs non-urban; ΔAIC = temperature model AIC minus urban model AIC; FDR-corrected p-values in parentheses.

| Trait | Model | Effect | numDF | denDF | *F* | *p* |
|---|---|---|---|---|---|---|
| Postdiscal NIR-r | Urban vs non-urban | Urban vs non-urban | 1 | 13 | 0.09 | 0.77 (0.89) |
| | | Region | 2 | 13 | 0.41 | 0.67 (0.99) |
| | | Sex | 1 | 284 | 16.84 | < 0.001 (< 0.001) |
| | | UvNU:Region | 2 | 13 | 1.84 | 0.20 (0.46) |
| | | UvNU:Sex | 1 | 284 | 6.18 | 0.01 (0.09) |
| | Temperature gradient | Temperature | 1 | 13 | 0.01 | 0.91 (0.91) |
| | | Region | 2 | 13 | 0.45 | 0.65 (0.92) |
| | | Sex | 1 | 284 | 15.93 | < 0.001 (< 0.001) |
| | | Temperature:Region | 2 | 13 | 1.18 | 0.34 (0.56) |
| | | Temperature:Sex | 1 | 284 | 1.88 | 0.17 (0.83) |
| Model comparison: Urban vs non-urban AIC = 2156.8, Temperature AIC = 2167.8, ΔAIC = 11 | | | | | | |
| Postdiscal Vis-r | Urban vs non-urban | Urban vs non-urban | 1 | 13 | 0.02 | 0.89 (0.89) |
| | | Region | 2 | 13 | 0.15 | 0.86 (0.99) |
| | | Sex | 1 | 284 | 163.01 | < 0.001 (< 0.001) |
| | | UvNU:Region | 2 | 13 | 1.18 | 0.34 (0.59) |
| | | UvNU:Sex | 1 | 284 | 3.33 | 0.07 (0.24) |
| | Temperature gradient | Temperature | 1 | 13 | 0.19 | 0.67 (0.88) |
| | | Region | 2 | 13 | 0.56 | 0.59 (0.92) |
| | | Sex | 1 | 284 | 163.26 | < 0.001 (< 0.001) |
| | | Temperature:Region | 2 | 13 | 0.97 | 0.40 (0.56) |
| | | Temperature:Sex | 1 | 284 | 0.14 | 0.71 (0.83) |
| Model comparison: Urban vs non-urban AIC = 2033.0, Temperature AIC = 2042.7, ΔAIC = 9.7 | | | | | | |

| Trait | Model | Effect | numDF | denDF | *F* | *p* |
|---|---|---|---|---|---|---|
| Wing tip NIR-r | Urban vs non-urban | Urban vs non-urban | 1 | 13 | 0.23 | 0.64 (0.89) |
| | | Region | 2 | 13 | 0.01 | 0.99 (0.99) |
| | | Sex | 1 | 283 | 26.88 | < 0.001 (< 0.001) |
| | | UvNU:Region | 2 | 13 | 2.88 | 0.09 (0.46) |
| | | UvNU:Sex | 1 | 283 | 1.41 | 0.24 (0.55) |
| | Temperature gradient | Temperature | 1 | 13 | 0.54 | 0.48 (0.88) |
| | | Region | 2 | 13 | 0.50 | 0.62 (0.92) |
| | | Sex | 1 | 283 | 26.71 | < 0.001 (< 0.001) |
| | | Temperature:Region | 2 | 13 | 1.87 | 0.19 (0.56) |
| | | Temperature:Sex | 1 | 283 | 0.28 | 0.59 (0.83) |
| Model comparison: Urban vs non-urban AIC = 2076.3, Temperature AIC = 2084.8, ΔAIC = 8.5 | | | | | | |
| Basal FW NIR-r | Urban vs non-urban | Urban vs non-urban | 1 | 13 | 0.46 | 0.51 (0.89) |
| | | Region | 2 | 13 | 0.47 | 0.64 (0.99) |
| | | Sex | 1 | 285 | 0.48 | 0.49 (0.57) |
| | | UvNU:Region | 2 | 13 | 2.23 | 0.15 (0.46) |
| | | UvNU:Sex | 1 | 285 | 0.63 | 0.43 (0.65) |
| | Temperature gradient | Temperature | 1 | 13 | 0.22 | 0.65 (0.88) |
| | | Region | 2 | 13 | 0.24 | 0.79 (0.92) |
| | | Sex | 1 | 285 | 0.58 | 0.45 (0.52) |
| | | Temperature:Region | 2 | 13 | 1.35 | 0.29 (0.56) |
| | | Temperature:Sex | 1 | 285 | 0.53 | 0.47 (0.83) |
| Model comparison: Urban vs non-urban AIC = 2398.8, Temperature AIC = 2406.9, ΔAIC = 8.1 | | | | | | |

| Trait | Model | Effect | numDF | denDF | *F* | *p* |
|---|---|---|---|---|---|---|
| Basal HW dorsal NIR-r | Urban vs non-urban | Urban vs non-urban | 1 | 13 | 1.87 | 0.19 (0.89) |
| | | Region | 2 | 13 | 0.36 | 0.71 (0.99) |
| | | Sex | 1 | 284 | 2.25 | 0.13 (0.19) |
| | | UvNU:Region | 2 | 13 | 0.88 | 0.44 (0.62) |
| | | UvNU:Sex | 1 | 284 | 0.54 | 0.46 (0.65) |
| | Temperature gradient | Temperature | 1 | 13 | 2.99 | 0.11 (0.75) |
| | | Region | 2 | 13 | 1.69 | 0.22 (0.92) |
| | | Sex | 1 | 284 | 2.31 | 0.13 (0.18) |
| | | Temperature:Region | 2 | 13 | 1.83 | 0.20 (0.56) |
| | | Temperature:Sex | 1 | 284 | 0.34 | 0.56 (0.83) |
| Model comparison: Urban vs non-urban AIC = 2418.0, Temperature AIC = 2423.1, ΔAIC = 5.1 | | | | | | |
| Basal HW ventral NIR-r | Urban vs non-urban | Urban vs non-urban | 1 | 13 | 1.41 | 0.26 (0.89) |
| | | Region | 2 | 13 | 0.53 | 0.60 (0.99) |
| | | Sex | 1 | 285 | 0.07 | 0.80 (0.80) |
| | | UvNU:Region | 2 | 13 | 0.40 | 0.68 (0.69) |
| | | UvNU:Sex | 1 | 285 | 0.20 | 0.66 (0.66) |
| | Temperature gradient | Temperature | 1 | 13 | 0.49 | 0.50 (0.88) |
| | | Region | 2 | 13 | 0.34 | 0.72 (0.92) |
| | | Sex | 1 | 285 | 0.05 | 0.82 (0.82) |
| | | Temperature:Region | 2 | 13 | 0.19 | 0.83 (0.83) |
| | | Temperature:Sex | 1 | 285 | 0.23 | 0.63 (0.83) |
| Model comparison: Urban vs non-urban AIC = 2317.9, Temperature AIC = 2325.4, ΔAIC = 7.5 | | | | | | |

| Trait | Model | Effect | numDF | denDF | *F* | *p* |
|---|---|---|---|---|---|---|
| Marginal HW NIR-r | Urban vs non-urban | Urban vs non-urban | 1 | 13 | 0.12 | 0.74 (0.89) |
| | | Region | 2 | 13 | 0.15 | 0.87 (0.99) |
| | | Sex | 1 | 280 | 2.58 | 0.11 (0.19) |
| | | UvNU:Region | 2 | 13 | 0.38 | 0.69 (0.69) |
| | | UvNU:Sex | 1 | 280 | 0.25 | 0.62 (0.66) |
| | Temperature gradient | Temperature | 1 | 13 | 0.10 | 0.75 (0.88) |
| | | Region | 2 | 13 | 0.03 | 0.97 (0.97) |
| | | Sex | 1 | 280 | 2.57 | 0.11 (0.18) |
| | | Temperature:Region | 2 | 13 | 0.73 | 0.50 (0.59) |
| | | Temperature:Sex | 1 | 280 | 0.01 | 0.94 (0.94) |
| Model comparison: Urban vs non-urban AIC = 2303.2, Temperature AIC = 2309.3, ΔAIC = 6.1 | | | | | | |

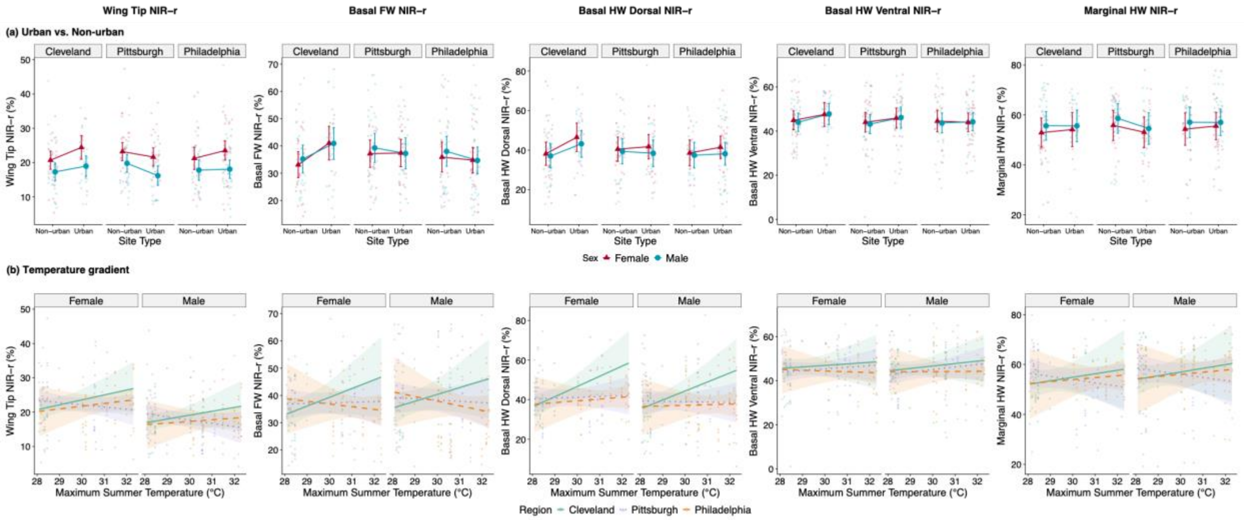

**Figure S4.** Effects of urbanization and temperature on near-infrared reflectance (NIR-r) of *Pieris rapae* at multiple wing regions on the forewing (FW) and hindwing (HW). Raw data are shown with 95% confidence intervals on predicted marginal means and estimated slopes. (A) Urban vs non-urban effects, shown separately for each region. (B) Continuous temperature gradient, shown separately for each sex.
